# Stiffness as a control factor for object manipulation

**DOI:** 10.1101/339101

**Authors:** Scott D. Kennedy, Andrew B. Schwartz

## Abstract

We act on the world by producing forces that move objects. During manipulation, force is exerted with the expectation that an object will move in an intended manner. This prediction is a learned coordination between force and displacement. Mechanically, *impedance* is a way to describe this coordination. As an efficient control strategy, object interaction could be anticipated by setting impedance before the hand moves the object. We examined this possibility with a paradigm in which subjects moved a handle to a specific target position along a track. The handle was locked in place until the subject exerted enough force to cross a specific threshold; then the handle was abruptly released and could move along the track. We hypothesized that this ballistic-release task would encourage subjects to modify their arm impedance in anticipation of the upcoming movement. If we consider the handle as an object, this paradigm loosely approximates the uncertainty encountered at the end of a reach when contacting a fixed object. We found that one component of arm impedance, stiffness, varied in a way that matched the behavioral demands of the task and we were able to dissociate stiffness from changes in force and displacement. We also found separate components of muscle activity that corresponded to stiffness and to changes in force. Our results show that subjects used a robust and efficient strategy to coordinate force and displacement by modulating muscle activity in a way that was behaviorally relevant in the task.

**New & Noteworthy:** The arm can behave like a spring, suggesting the concept of exerting force to move an object by selecting a spring of a certain length and stiffness that, respectively, depend on the movement and force requirements of the task. We show that these spring-like characteristics describe the strategy used to arrest a pre-loaded handle. These results extend our understanding of the arm’s spring-like behavior to include force and movement constraints, important factors for object interaction.

## Introduction

Manipulating objects is fundamental to human behavior and requires flexible, coordinated control of both force and movement (Kawato 1999; Wolpert and Ghahramani 2000; Flanagan et al. 2006; Franklin and Wolpert 2011). Identifying a control scheme that takes place during this behavior would be a step toward detecting and understanding the brain signaling underlying the way we interact with objects. Depending on the task conditions, multiple strategies can be used to achieve this coordination. The most challenging behaviors are those in which rapid, precise manipulation takes place. While humans perform these movements with great skill, the control principles underlying this behavior is a topic of interest for both scientists and engineers.

Roboticists use control schemes that utilize rapid feedback to monitor ongoing changes in displacement and force. To manipulate an object, motion of the robotic effector is controlled precisely by exerting the force needed to achieve the movement. In real-world conditions, this can be problematic when unexpected collisions take place or if interaction with the object leads to large changes in force -- for instance, when displacement is measured inaccurately. The generation of these large forces makes robots dangerous in the workplace, less than ideal for tasks that rely on rapid and precise object interaction, and renders robots unsuitable for interaction with humans.

In contrast, human manipulative behavior can be complex, precise and rapid-despite noisy sensory information, long feedback delays and muscles with slow, nonlinear dynamics. How this takes place despite these biological constraints is an open question. For slow movements, a feedback strategy could be used to continuously monitor the object’s movement and to adjust the exerted force (Kalaska and Crammond 1992; Scott 2004; Scott et al. 2015). If interaction with the object is predictable, then fast movements could be performed using a feedforward strategy to plan the time-varying forces that produce the desired movement (Kawato 1999). However, there is always some amount of uncertainty when interacting with an object (Shadmehr and Mussa-Ivaldi 1994; Burdet et al. 2001; Rancourt and Hogan 2001; Takahashi et al. 2001; Milner and Franklin 2005a).

Modulating the arm’s mechanical impedance by coordinating force and kinematics has been proposed as a strategy for handling uncertain interaction dynamics between the hand and external forces (Bizzi et al. 1982, 1984; Hogan 1984a; Flash and Hogan 1985). Mechanical impedance is the force which opposes changes in movement, i.e., position, velocity, acceleration, etc. (Hogan 1984b, 1985a; Mussa-Ivaldi et al. 1985). At least three aspects of arm impedance, operating at different time scales, are relevant for object interaction. First, the current state of the musculotendon tissue, specified by ongoing neural activity, exerts force instantaneously to impede changes in position (Hill 1950). Second, spinal reflex loops modulate muscle force to impede stretch with latencies of tens of milliseconds (Joyce and Rack 1969; Rack and Westbury 1974; Nichols and Houk 1976; Pruszynski et al. 2009; Crevecoeur and Scott 2014). Finally, cortical reflex loops engage cortical networks at longer latencies but may still operate in a way that is covert to the subject (Kurtzer et al. 2009; Dimitriou et al. 2013; Pruszynski et al. 2014). The modulation of musculotendon tissue properties by reflex loops could be integral components of a feedback control strategy to counteract uncertainties encountered during object manipulation. A compliant arm and hand that yields predictably upon object interaction may minimize the need for moment-by-moment updates to a control signal. This type of control is likely a contributing factor to the fast and robust movements characteristic of human manipulation.

Arm impedance has been studied in a variety of experimental conditions. In posture-control paradigms, subjects held a manipulandum at an equilibrium position to resist randomly imposed displacements (Mussa-Ivaldi et al. 1985). When instructed to “resist” or “not resist” the displacements, subjects modulated their arm impedance to generate the required force needed to return to the equilibrium position (Lacquaniti et al. 1982). In addition, subjects could modulate arm impedance when instructed to co-activate different groups of antagonist muscles (Gomi and Osu 1998; Osu and Gomi 1999). However, it is unclear how the results of these studies can be extrapolated to real-world movements in which both posture and the equilibrium position change during object manipulation (Gomi and Kawato 1997; Darainy et al. 2007).

A variation of the posture-control paradigm required subjects to exert an isometric force. Under this paradigm, it was found that subjects adopted an arm impedance that was proportional to force (McIntyre et al. 1996). Again, different values of arm impedance could be achieved by co-activating different groups of antagonist muscles (Gomi and Osu 1998), but the range of this modulation was constrained by the level of isometric force (Perreault et al. 2002).

In addition to posture maintenance, arm impedance can also be used to constrain the arm spatially along an equilibrium trajectory as it moves toward a target (Bizzi et al. 1982, 1984; Hogan 1984a; Flash and Hogan 1985). A path toward the target would consist of a series of equilibrium positions. An arm following this trajectory would be resistant to perturbations away from these positions. If the equilibria were specified sequentially, the arm would be propelled toward the target by the continuous change in impedance. In this way, the control strategy could be modeled as a spring-mass-damper (stiffness-inertia-damping) that moves along the equilibrium trajectory, exerting force to pull the object along behind it. In subsequent studies testing this hypothesis, subjects were instructed to relax their arms as much as possible and although it was found that arm impedance varied during movement, the equilibrium trajectories were complex (Gomi and Kawato 1996, 1997). In later studies, subjects moved their arms through unstable force fields with varying amounts of uncertainty (Takahashi et al. 2001; Perreault et al. 2002; Osu et al. 2003; Milner and Franklin 2005b). Again, the subjects were able to modulate the impedance of their arms to complete the movements successfully. The finding that impedance could be steered to compensate for directionally-specific instability (Burdet et al. 2001; Franklin et al. 2007; Kadiallah et al. 2011) was taken as evidence for an explicit impedance controller with an internal model of environmental instability (Franklin et al. 2007).

In general, internal models are invoked to explain how predictions can be made to account for the inherent delays of the motor system. In particular, setting a value of impedance in anticipation of object interaction could be useful in minimizing the effect of these delays. Evidence that subjects change their impedance during a task to anticipate object contact was found using a task in which subjects caught a falling ball (Lacquaniti et al. 1993). Pseudorandom force pulses were used to measure arm impedance throughout the task and it was found that subjects changed the direction and magnitude of that impedance shortly before the ball made contact with the hand.

In order to further characterize anticipatory changes in impedance, we designed a ballistic-release paradigm that required subjects to pull on a handle with different levels of force and to move the handle to various target positions. We found that arm impedance varied with task conditions in a way that could be separated from its linkage to force. In addition, arm impedance and force were related to separate components of muscle activity. Not only do these results support the conclusions of previous studies showing that impedance is controlled explicitly, they suggest that it may be specified before an expected object interaction takes place. The preset impedance allows the forces following the perturbation to direct the arm and hand to an intended target.

## Materials and Methods

### Subjects

Five subjects -- four men and one woman between the ages of 20 and 40 -- performed a ballistic-release task with their right arm. All subjects were right handed, had no known neurological deficits, and gave written and informed consent to participate in the study. The protocol was approved by the University of Pittsburgh’s Institutional Review Board.

### Experimental design

The objective of our study was to determine how task constraints modulate the anticipatory changes in arm impedance and, more specifically, how these constraints modulate the relation between arm impedance and force. In the task, subjects were required to overcome four force thresholds and to arrest their movement in four different target zones. In theory, the arm could move to the correct target zone if the arm’s equilibrium position was adjusted so that the force would be zero when the handle was in the target zone (Feldman 1966, 1986; Polit and Bizzi 1979). For the same target zone and equilibrium position, the force of the arm pulling on the handle could be increased by increasing arm impedance. Thus, the equilibrium position and arm impedance could be preset for a given target zone and force threshold, eliminating the need to rely on corrections during the movement. Instead of emphasizing corrective movements, this simple paradigm focuses on the anticipatory control that is likely to take place during object manipulation.

### Behavioral paradigm

Subjects were seated with their torsos restrained to minimize extraneous movement during the task. A linear manipulandum was mounted approximately 35 cm in front of the subject at shoulder height (Figure 1A). The manipulandum consisted of a handle mounted on a sled that could be moved along a straight track. The device was oriented in the frontal plane with the starting end of the track aligned to the subject’s left shoulder. An electromagnet (Rectangular, 12V DC, 8W, McMaster-Carr, Chicago, IL) was activated to lock the sled in place with a microcontroller (Arduino Mega 2560) using custom software (dragonfly-msg.org). A start button was in line with the track, 15 cm to the right of the subject’s right shoulder (Figure 1A).

**Figure 1.**
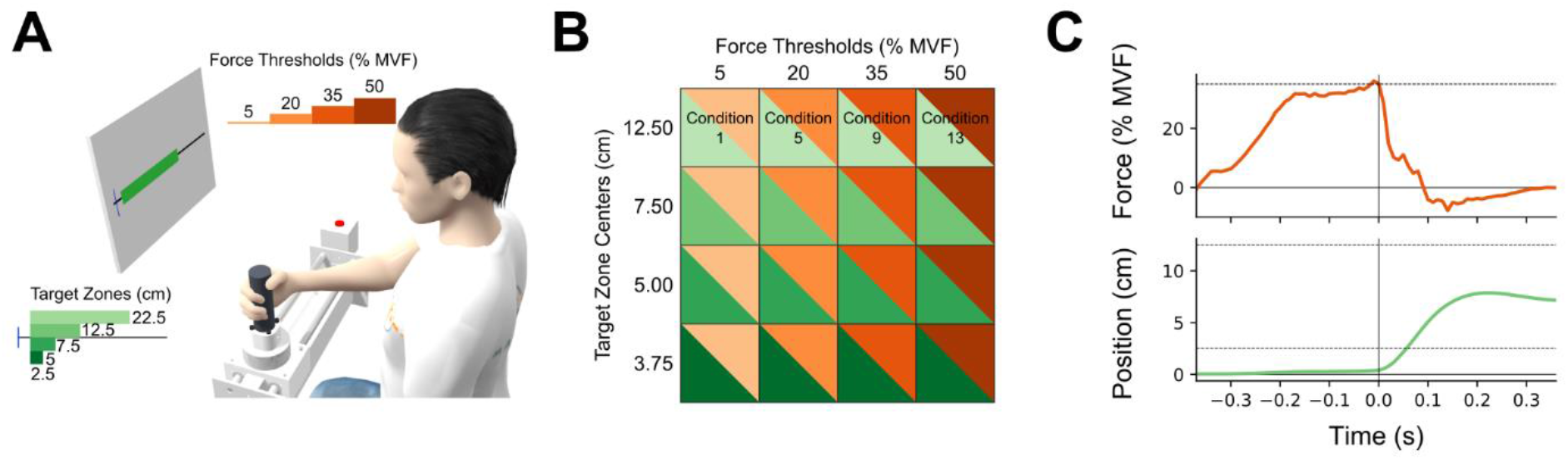
Each trial required the subject to control both force and movement. **(A)** The subject was seated in front of a handle with the torso restrained. A monitor displayed the position of the handle along a track (vertical blue line) and a target zone (green rectangle). To initiate a trial, the subject first pressed the start button (to the right of the track) and then grasped the handle. The subject pulled on the handle while it was locked in place until a force threshold was crossed (MVF, maximal voluntary force). The handle was then unlocked to move freely along the track and the subject had to stop and hold the handle within the target zone for 300 ms. **(B)** A single task condition was composed of a force threshold and a target zone. The subject completed 20 trials per condition, beginning with the lowest force threshold/farthest target zone and ending with the highest force threshold/closest target zone. In this way, every combination of force threshold and target zone was systematically sampled. The target zone and force threshold colors will be used throughout the Figures. **(C)** Time-series of force and displacement were measured for each trial (a single representative trial is displayed here). When the subject pulled on the locked handle, the force increased isometrically. At time 0, the handle was unlocked, the force decreased rapidly, and the handle’s position moved to the target zone and then stopped within it. The dashed line on the force plot was the force threshold and the pair of dashed lines on the position plot were the near and far boundaries of the target zone.

Each subject was instructed to (1) use the right hand to press the button and then reach to grasp the handle, (2) pull on the handle with enough force to unlock it, and (3) position the handle within a specified target zone. Real-time feedback about the handle’s location and the target zone was displayed on a monitor in front of the subject. However, the subject did not receive any direct feedback of the pulling force exerted on the handle or of the force necessary to unlock the handle.

To be successful, the subject needed to pull with enough force to unlock the handle and hold it in the specified target zone for 300 ms. Exiting the target zone before 300 ms had elapsed caused the trial to fail. An auditory cue indicated success or failure. The subject then returned the handle to the lock position and initiated the next trial by pressing the start button. Successful and unsuccessful trials were both included in our analyses.

Subjects were given 6 seconds to unlock the handle. The trial was aborted if the subject failed to move the handle in this period and the trial was reinitiated by pressing the start button. These incidents were not counted as a failure nor included in the analyses.

#### Force thresholds

Four force thresholds were chosen as a percentage of the subject’s maximum voluntary force (MVF). At the beginning of the experimental session, each subject was asked to grasp the handle and pull as hard as possible while the handle was locked in place. We measured the maximum force along the direction of movement and set the four force thresholds at 5, 20, 35, and 50% of the MVF (Figure 1A and 1B).

To overcome the force threshold, only force along the movement direction was considered. Because of hardware limitations, there was an unavoidable random 10-30 ms delay between threshold detection and magnet release. This delay was not visible to the recording system, leading to the decision to use movement onset as an alignment time (see *Behavior alignment*).

#### Target zones

The target zones were displayed on a monitor at eye-level in front of the subject (Figure 1A). All four target zones began 2.5 cm from the handle’s lock position, measured along the track. The end of the four targets were 5, 7.5, 12.5, and 22.5 cm from the handle’s lock position. Target-zone centers were located 3.75, 5, 7.5, and 12.5 cm from the start position (Figure 1B). The target was shown as soon as the start button was pressed.

A blue bar represented the handle’s real-time position on the monitor. The handle’s position was measured by a microcontroller (Arduino Mega 2560) that sampled a linear potentiometer (SoftPot 300.00mm, Spectra Symbol) at 100 Hz. Voltage output of the linear potentiometer changed as a wiper, attached to the sled, moved along the potentiometer’s surface.

We wanted to encourage the subjects to modulate arm impedance. The target zones were chosen to have different widths because arm impedance has been shown to increase with positional accuracy (Gribble et al. 2003; Selen et al. 2006). Here we label target zones by their center positions.

#### Task conditions

A single task condition was composed of a force threshold and a target zone. There were four force thresholds and four target zones, resulting in 16 task conditions (Figure 1B).

The task conditions were presented in blocks of 20 repeated trials. The order in which the task conditions were presented remained the same for each subject, beginning with the lowest threshold and farthest target. This block was followed by another with the same threshold, but with the second-farthest target. Two more blocks were completed with targets that moved progressively closer to the handle’s lock position. The next lowest threshold was then presented for the farthest target and the pattern continued. The last task condition was the highest threshold and the closest target.

The subject was able to rest whenever necessary to prevent fatigue. In addition, a 30-second rest period was given between task conditions. A longer 60-second rest was given between task conditions when the threshold changed, e.g., when the target reset from the closest target to the farthest target. On average, the total time from one start button press to the next was approximately 4.5 seconds.

### Signal measurement

During the task, we measured three signals: (1) the force of the subject pulling on the handle; (2) the surface electromyographic (EMG) activity of 8 muscle groups; and (3) the 3D position of optical markers placed on the handle and the subject’s hand, lower arm, upper arm, and torso.

#### Force data

Force was measured using a six degree-of-freedom force transducer (Delta FT, ATI Industrial Automation, Inc.) and sampled at 100 Hz. The transducer was mounted between the handle and the sled. We refer to force as the linear force exerted by the hand on the handle in the direction co-linear with the track.

#### EMG data

The surface muscle electromyogram (AMT-8, Bortec Biomedical) was collected at 2000 Hz (ADLink DAQ-2208 data acquisition card; ADLink Technology Inc.). Eight differential signals were recorded using sixteen electrodes (Pediatric electrodes, Vermed Inc.). Two electrodes were placed over the following muscle groups: wrist flexors, wrist extensors, elbow flexors, elbow extensors, anterior deltoid, posterior deltoid, pectoralis major, and rotator cuff. The electrodes were placed using anatomical landmarks and verified by displaying the signal on an oscilloscope and asking the subject to activate the different muscle groups. A global reference electrode was placed on the back of the right hand.

#### Motion tracking

Movement was measured using a passive, infrared motion tracking system (Nexus 1.8.5, Vicon, Inc.) that sampled at 100 Hz. The system consisted of twelve cameras and eighteen markers that were 10 mm in diameter. One marker was placed on the handle. The remaining markers were placed on the subject.

Four markers were on the subject’s torso at the right acromion, left acromion, jugular notch, and xyphoid process. Two markers were placed on the elbow joint at the medial and lateral condyles of the humerus and two markers were placed on the wrist joint at the radial and ulnar styloid processes. A set of three markers were placed as a rigid triangle on the lateral side of the subject’s upper arm, another set on the lateral side of the subject’s lower arm, and the final set on the dorsal side of the subject’s hand.

Although the handle’s real-time position was displayed to the subject using the signal from the linear potentiometer, this signal was noisy, so the marker on the handle was used for analysis.

### Data preprocessing

#### EMG

For each muscle, the raw differential EMG signal was high-pass filtered at 100 Hz. It was then mean centered and scaled by the standard deviation. Finally, the signal was squared, low-pass filtered at 30 Hz, and square-root transformed. The resulting signal envelope was down-sampled to 100 Hz.

#### Motion tracking

The motion tracking markers were manually labelled offline. Gaps in each marker’s trajectory were filled using spline or source fitting tools available in the motion tracking software.

To define the reference frame of the track, we performed principal component analysis on the handle’s 3-D position in the motion tracking reference frame. The first component was the direction along the track and the remaining vectors were ordered so that the second dimension pointed away from the subject and the third dimension pointed upwards. The origin of the reference frame was the handle’s start position.

Velocity and acceleration were calculated from the trial-averaged position using successive application of a Savitzky-Golay filter with a window length of 7 time bins, a polynomial order of 3, and a derivative order of 1 (scipy.signal.savgol_filter).

#### Joint angles

Joint angles were calculated using musculoskeletal modeling software (OpenSim) (Delp et al. 2007) and a generic musculoskeletal model of a human torso (Holzbaur et al. 2005). The model was scaled using the markers placed on bony landmarks. Joint angles were found using OpenSim’s inverse kinematics algorithm. In brief, the algorithm found the joint angles that minimized the error between the measured marker positions and a set of virtual marker positions placed on the model. Joint centers were calculated using the joint angles and the scaled model.

### Behavior alignment

#### Force ramp

Each trial included a time period when the subjects isometrically increased the force they exerted on the handle, which we refer to as the force ramp. The force ramp began when the force exerted on the handle rose above 1 N for the last time before the force threshold was crossed and ended at movement onset. Movement onset was determined as the first time, after the force threshold was crossed, when velocity rose above 10% of the maximum velocity for that trial.

The duration of the force ramp varied primarily as a function of the force threshold, with lower thresholds having a shorter duration than longer thresholds. To compare task conditions with the same force threshold and different target zones, we averaged the force ramp duration across trials with the same force threshold and scaled the duration of each trial’s force ramp to match the average using a piecewise cubic hermite interpolating polynomial. The trial-averaged data during the force ramp were then used in the analyses.

#### Movement

Behavior following movement onset was not scaled in time because of the importance of the position time derivatives in the impedance analysis. Instead, the trial-averaged behavior is presented for a consistent 500 ms window across all task conditions, beginning at movement onset. The impedance analysis was performed on a 200 ms time window, beginning at movement onset.

The impedance hypothesis predicts that the equilibrium position is where the handle position will be stopped and held, making it an important factor in our stiffness analyses. Although subjects were instructed to hold the handle within the target zone for 300 ms, it was common for subjects to return the handle to the start position without holding when the handle prematurely exited the target zone (unsuccessful trials). In addition, the wide target zones made it possible for the subject to enter the target zone and begin to return the handle to the start position while still remaining in the target zone for 300 ms (a successful trial). Because both successful and unsuccessful trials were included in our analyses, these unanticipated behaviors made it difficult to consistently evaluate the effect of the estimated equilibrium position across all task conditions.

To address this difficulty, we examined the position trajectory when the subjects did hold the handle at a final position and found that the time when velocity first changed sign, which we call the arrest position, was a close approximation to the hold position. When the handle asymptotically approached the hold position, the arrest position was nearly identical to the hold position. When the handle oscillated slightly toward the end of the movement, the arrest position was slightly farther from the start position than the hold position. Therefore, the arrest position was used to evaluate the estimated equilibrium position and to determine the effect of movement constraints on the estimated stiffness.

### Physical dynamical model

During the movement, the arm was modeled as a physical dynamical system consisting of four impedance components. Each of these components exerted force as a function of movement according to the following equation,

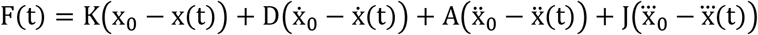

where F is the force that the arm exerts on the handle; x, ẋ, ẍ, 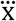 are the position, velocity, acceleration, and jerk of the hand; x_0_, 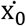, ẋ_0_, 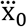 are the equilibrium position, velocity, acceleration and jerk; and K, D, A, J are the arm impedance elements.

We assumed that 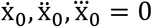, resulting in five free parameters: x_0_, K, D, A, J.

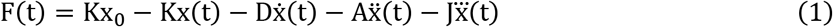

These parameters were found by fitting equation 1 to the trial-averaged force and movement during a 200 ms time window, beginning at movement onset, for each of the 16 conditions using iterative least-squares optimization. The mean and 95% confidence interval of the model parameters were found by resampling the trial data 1000 times, with replacement, from the 20 trials per condition. For each resample, we averaged the 20 trials per condition and fit the model parameters.

### EMG analysis

We were interested in characterizing the EMG pattern that correlated with arm stiffness, which can increase when antagonist muscle groups co-activate. When this is the case, EMG can change without a change in the force exerted on the handle. In our analysis, we distinguished between changes in the EMG pattern that were most correlated with changes in force (potent EMG activity) from those patterns that were less correlated (null EMG activity).

#### Regressing force on EMG

We regressed force on EMG during the force ramp using the trial-averaged force and EMG signals from the 16 task conditions. We first fit the linear model according to equation 2

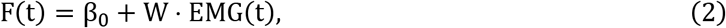

where F is force along the track, β_0_ is the offset, W is the coefficient vector, and EMG is muscle activity from 8 muscle groups. We then used this model to predict force from EMG for each task condition.

#### Separating potent EMG and null EMG

We isolated the EMG pattern that most correlated with changes in force using singular value decomposition (SVD) (Kaufman et al. 2014) on the [1 × M] coefficient vector in equation 2,

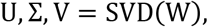

where V is an [M × M] rotation matrix, Σ is a [1 × M] scaling matrix, U is a [1 × 1] rotation matrix, and M is the number of muscle groups. EMG projected onto the first dimension (row) of V was most correlated with changes in force. We called this the potent EMG activity:

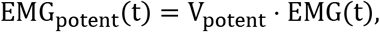

where V_potent_ is a [1 × M] vector that is the first row of V. If the relation between EMG and force was perfectly linear, then EMG projected onto the remaining 7 dimensions, i.e. rows 2 through M of V, would not be correlated with changes in force. We summarized this EMG activity by performing principal components analysis on the non-potent EMG and found the single dimension that captured the most variance. EMG was projected onto this dimension and the result was considered “null EMG”:

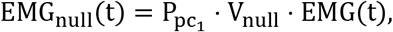

where V_null_ is an [M-1 × M] matrix that is rows 2 through M of V and P_pC_1__ is a [1 × M-1] eigenvector of V_null_ · EMG(t).

#### Testing the effect of arrest position on the regression of stiffness on force threshold

The partial F-test determines if additional parameters improve the explanatory power of a regression model. It employs two nested models, a full model and a restricted model. The restricted model consists of a subset of the parameters from the full model. The two models’ residual sum of squares are compared according to

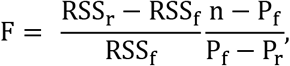

where F is the F-statistic, RSS_f_ and RSS_r_ are the residual sum of squares of the full and restricted model, n is the number of observations, and P_f_ and P_r_ are the number of parameters in the full and restricted model. A higher F-statistic indicates more explanatory power in the full model. The null hypothesis assumes a value of 0, indicating that the full model does not have more explanatory power than the restricted model. Statistical significance is tested using the F-distribution with (P_f_ − P_r_, n) degrees of freedom.

The partial F-test was used to test the hypothesis that adding the arrest position as a parameter would improve the regression of stiffness on force threshold. Equation 3 describes the full model and equation 4 describes the restricted model,

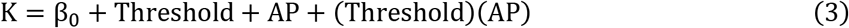

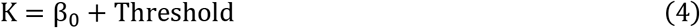

where, across all task conditions, K is stiffness, Threshold is the force threshold, AP is the arrest position, and β_0_ is the offset.

## Results

Five subjects performed a ballistic-release task that required them to pull on a handle with different levels of force, followed by movement to the different target zones. Successful trials were achieved by exerting enough force to unlock the handle while still controlling the subsequent movement. Below, we highlight the results from subject 1 and include the remaining subjects in the supplementary material. Summary statistics across all subjects are also presented.

### Force and movement varied with the four force thresholds and four target zones

The subject pulled on the handle with four levels of force and moved the handle to four target zones (Figure 2). Movement onset began at time 0 and, at this time, force for the same threshold was similar across targets (Figure 2A). At or near the end of the movement, 500 ms after movement onset, position was separated across targets for the same threshold (Figure 2B).

**Figure 2.**
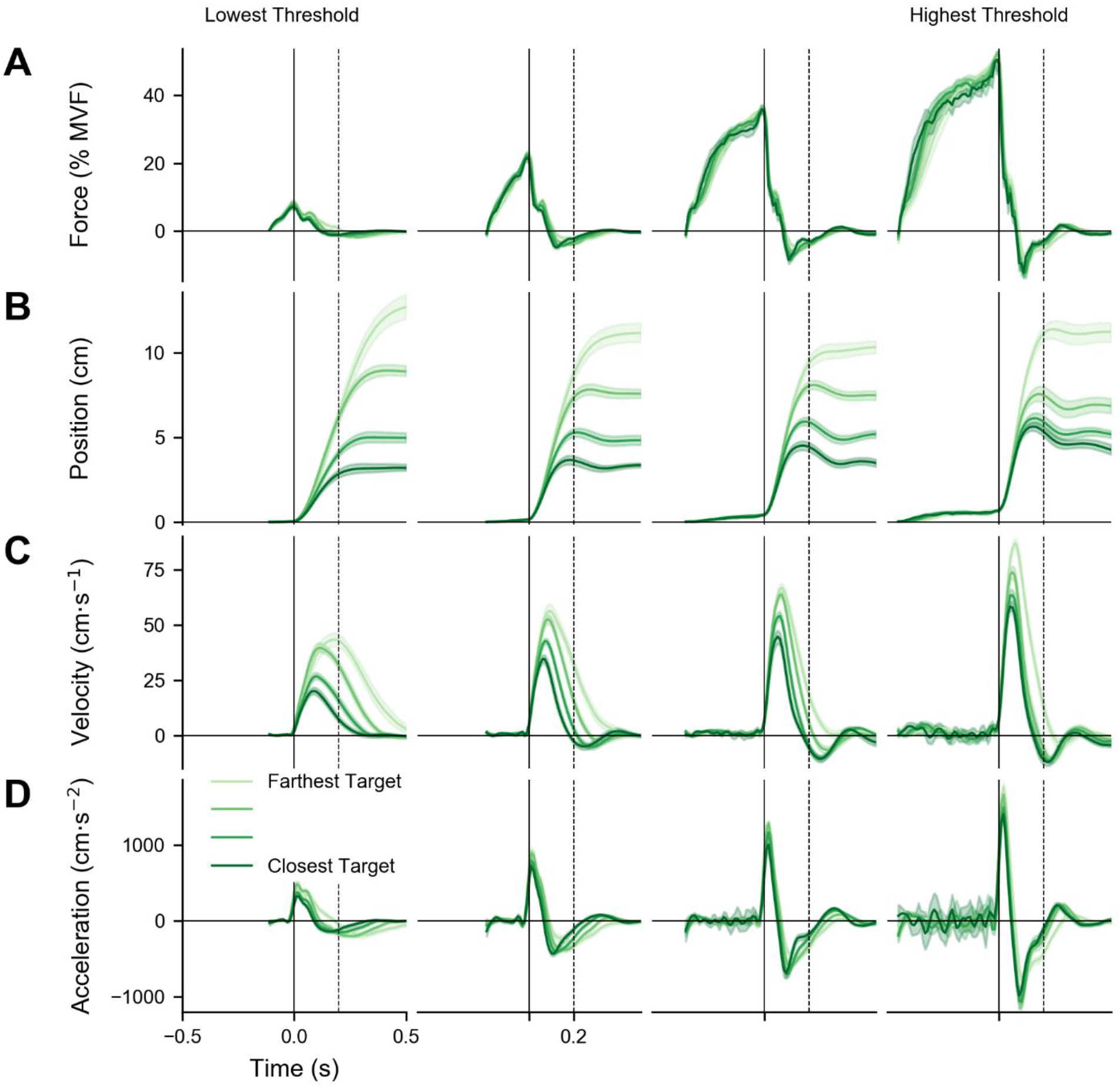
Behavioral results. The data are trial-averaged from one subject. Shading represents the median and 95% confidence interval. Each plot depicts four target zones with the same force threshold. Movement onset began at time 0 (solid vertical line). **(A)** Force varied across thresholds. For a given threshold, force was similar across targets. **(B)** Position varied across targets. Subjects tended to overshoot the target zone as the threshold increased and the target moved closer. **(C)** Velocity varied with target and threshold. The maximum velocity was related to both target and threshold. **(D)** Acceleration varied with threshold. For a given threshold, initial deceleration was similar across targets. Results for the remaining subjects are shown in Supplementary Figure 1.

For each signal and task condition, we found the median and 95% confidence interval of the trial-averaged data. We used a bootstrap to re-calculate the trial average 1,000 times. Each calculation used a random sample of the trials, with replacement.

For this subject, force increased rapidly and began to plateau as it approached the threshold. This pattern was more pronounced for higher thresholds and closer targets. The plateau suggests that the subject could approximate the force that would unlock the handle but was unable to accurately predict the exact timing of when the handle would be unlocked. After movement onset, the force decreased rapidly, falling below 0 to slow the handle and then approaching 0 as the movement of the handle reached a steady position. This trend was consistent across all subjects (Supplementary Figure 1). The MVF for this subject was 160 N.

For this subject, the movements were usually arrested around the center of the target, after some initial overshoot. The overshoot remained in the target zone, except for the closest target and the highest threshold combination, where it often extended beyond the target zone, reflecting the extreme difficulty of this task condition. The overshoot was more pronounced for closer targets and higher thresholds which corresponds to a physical dynamical system having various values of impedance.

Before movement onset, the position of the handle was fixed at 0, with some handle bending (on the order of millimeters) for high thresholds. Because all trials were included in the analysis, there was substantial across-subject variability in the positions of the handle at the end of the trial (Supplementary Figure 1). For a given subject and task condition, the across-trial variability of the positions was small, suggesting that the subject had selected a consistent movement strategy for that condition.

The timing of the maximum velocity was similar across all task conditions (Figure 2C). However, the magnitude of the maximum velocity depended on both the target zone and the force threshold, increasing for targets farther from the lock position and for higher thresholds.

As expected, the magnitude of the maximum acceleration depended on the force threshold (Figure 2D). The initial acceleration values are inaccurate due to the numerical differentiation of position. For a pre-loaded release, the initial acceleration should be nearly a step function and begin at time 0. The initial deceleration was similar across target zones with the same force threshold.

Task conditions varied from extremely easy to nearly impossible (Table 1, statistics across subjects). The first four task conditions began with the lowest threshold and farthest target (top left element in Table 1 and proceeding down). Success rates for these initial conditions were lower than expected because of some occasional confusion about the exact requirements of the task. However, after a few trials, the subjects moved quickly and smoothly (Figure 2B and Supplementary Figure 1), although the 95% confidence interval was slightly wider for the handle position during these first four task conditions (different targets with the lowest threshold) compared with the remaining the task conditions.

**Table 1.** Success rate for each task condition, mean +/− standard error across subjects. On successful trials, the subject moved the handle into the target zone and remained there for 300 ms. Exiting the target zone before 300 ms caused the trial to fail. The difficulty of the task generally increased as the force threshold increased and as the distance to the target zone decreased. The order of the task conditions started in the upper left element and proceeded down the column.

|  | Lowest Threshold |  |  | Highest Threshold |
| --- | --- | --- | --- | --- |
| Farthest Target | 91.4% +/- 2.4 | 100% +/- 0 | 100% +/- 0 | 100% +/- 0 |
|  | 97.0% +/- 1.8 | 99.0% +/- 0.9 | 99.0% +/- 0.9 | 94.0% +/- 3.6 |
|  | 82.4% +/- 7.6 | 95.0% +/- 2.0 | 77.3% +/- 10.8 | 54.0% +/- 16.5 |
| Closest Target | 71.0% +/- 15.4 | 68.3% +/- 7.8 | 39.0% +/- 14.4 | 5.0% +/- 3.5 |

### Arm impedance varied with force and movement

We modeled the arm as a physical dynamical system consisting of four elements with five free parameters. For each task condition, we averaged the force exerted on the handle and the handle’s movement across trials. We then used least-squares optimization to find the model parameters that minimized the difference between the actual and predicted force over 200 ms beginning at movement onset (see Methods, equation 1).

The equilibrium position (EP) of the physical model represents the position of the handle for which the arm would exert zero force. In the context of this task, the EP can be considered the position where the handle would come to rest and could reflect the intended hold position (Figure 3 and Supplementary Figure 2). The following describes the results from Subject 1. We observed that the distance to the EP increased for arrest positions farther from the handle’s lock position (Figure 3). However, the distance to the EP was consistently shorter than the distance to the arrest position. The difference between the arrest position and the EP is suggestive of an underdamped system. Although individual subjects sometimes arrested their movements at locations that were outside the specified target zones, the match between EP and the arrest position followed the general trend displayed by this subject (see also Supplementary Figure 2).

**Figure 3.**
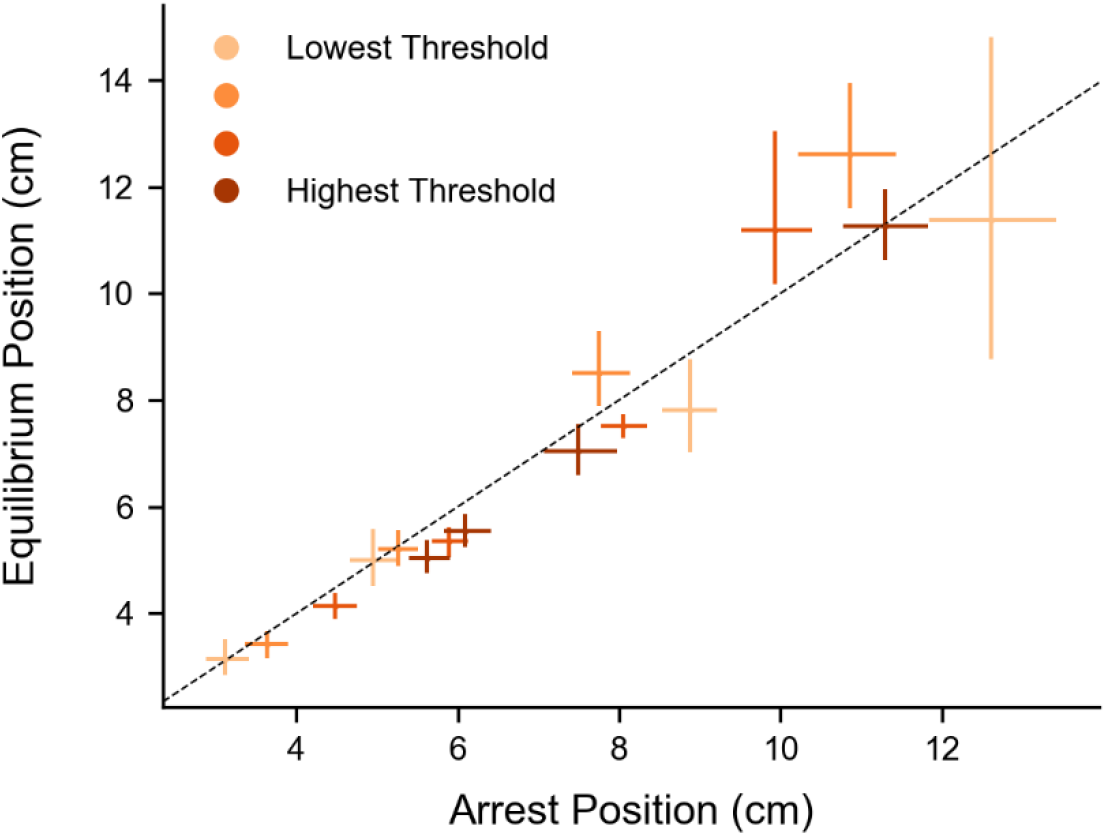
The equilibrium position (EP) co-varied with the arrest position. The distance to the EP increased as the distance to the arrest position increased. However, the distance to the EP was less than the distance to the arrest position. This pattern was consistent across all subjects (Supplementary Figure 2) and is suggestive of an underdamped system. Error bars indicate median and 95% confidence interval. Dashed line is the unity line. The model was fit using 200 ms of data beginning at movement onset.

The task was designed so that the same target was specified for different force thresholds. If the EP depended on the target, then the different force thresholds could be crossed by selecting different arm impedances. This would be an efficient control strategy for the ballistic-release task. We found that stiffness remained consistent across trial for each task condition (Supplementary Figure 3) and that stiffness increased with force threshold for a given target zone (Figure 4 and Supplementary Figure 4), consistent with previous results (McIntyre et al. 1996). Although this relation was approximately linear for a given target zone, across all task conditions there was distinct structure in the effect of target zone on the relation between stiffness and force threshold.

**Figure 4.**
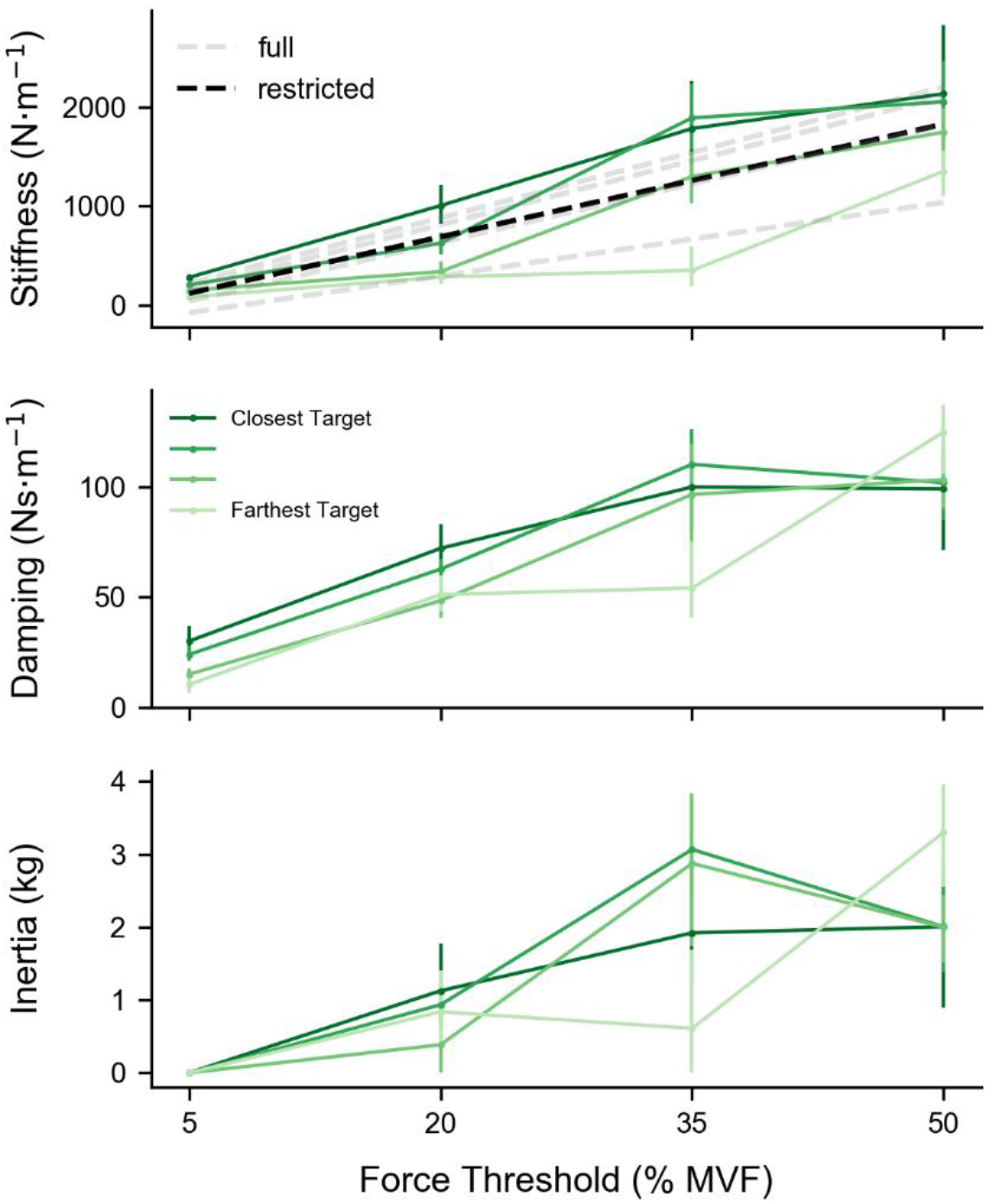
Stiffness co-varied with both force threshold and arrest position. The data are from subject 1. Although stiffness was proportional to threshold for a given target, across all task conditions the linear fit that included arrest position was better able to explain stiffness. The dashed grey lines represent the linear fit of the full regression of stiffness on force threshold and arrest position. The dashed black line represents the linear fit of the restricted regression of stiffness on force threshold alone. Damping and inertia increased with force threshold. However, there was not a strong relation between these impedance elements and the arrest position. Error bars indicate median and 95% confidence interval. The physical dynamical model was fit using 200 ms of data beginning at movement onset.

For each subject, we quantified the target-dependent structure by comparing the linear fit of two nested regression models using a partial F-test (Figure 5). The full model regressed stiffness on both threshold and arrest position (grey lines in Figure 4). The restricted model regressed stiffness on threshold alone (black lines in Figure 4). Adding the arrest position to the regression increased the predictive power as evidenced by the F-statistic’s p-value, which was below 0.05 for every subject: 0.003, 0.002, 0.008, 0.03, 0.0006; F-distribution with (2, 12) degrees of freedom.

**Figure 5.**
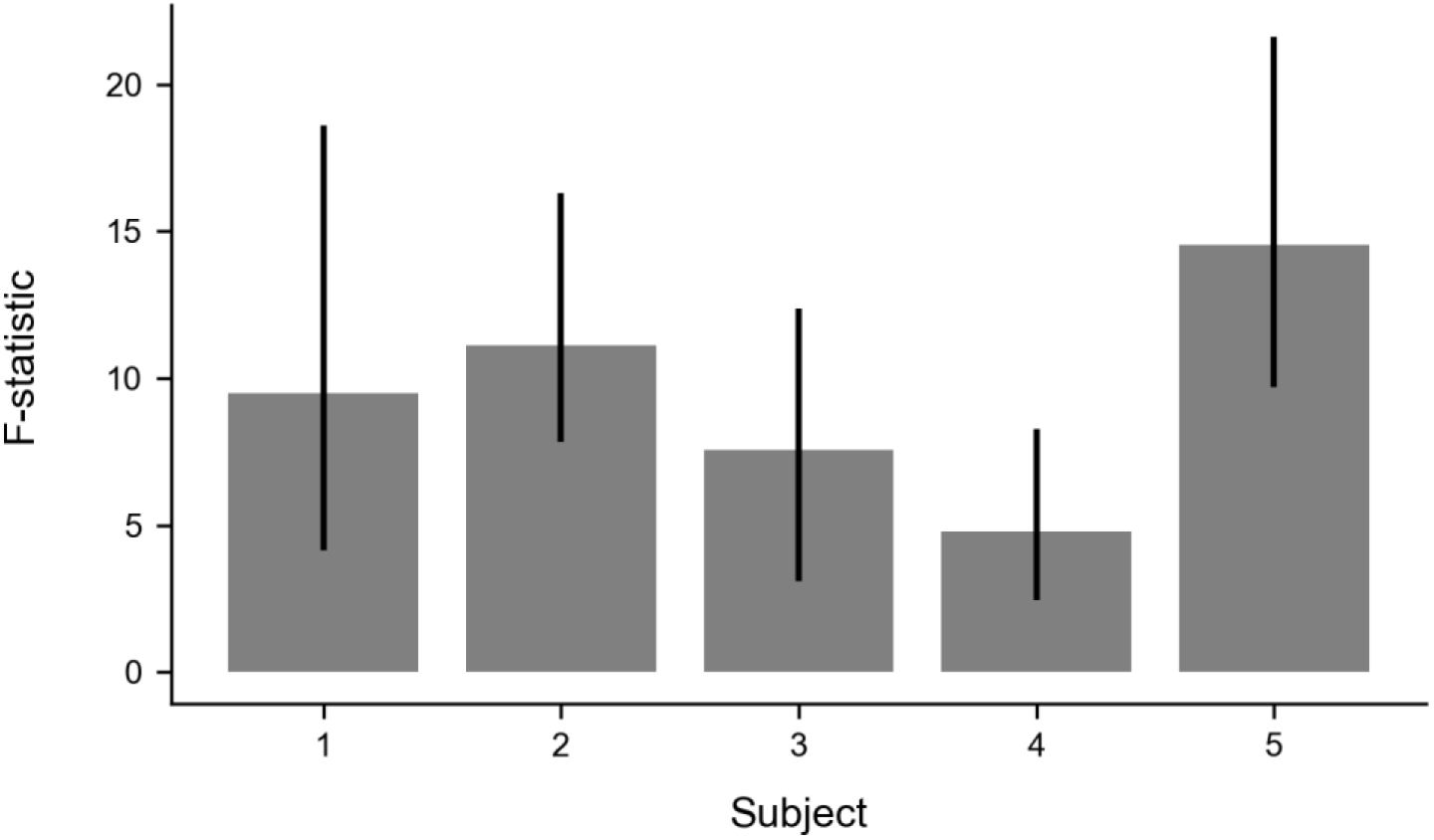
Stiffness variability is best explained by both force and movement information. F-statistic from a partial F-test comparing the regression of stiffness on force threshold with and without additional arrest position information. The additional arrest position information improved the stiffness regression for all subjects, with subject 1 and 5 showing the largest effect. Error bars represent median and 95% confidence interval.

Although damping and inertia generally increased with force threshold, there was not a consistent relation between these impedance elements and target zone (Figure 4). When stiffness increases more than damping, the physical dynamical system can become underdamped and tend to oscillate, which may explain the target overshoot and movement oscillations seen in Figure 2.

### Arm posture before movement did not vary across task conditions

Although we suspected that subjects might vary their arm posture for different task conditions (Trumbower et al. 2009), we found the arm’s configuration to be remarkably consistent. For subject 1, the deviation of the joint centers during the 200 ms before movement onset across all task conditions were as follows: torso 0.98 cm [0.21, 1.99], shoulder 1.23 cm [0.33, 2.39], elbow 1.1 cm [0.27, 2.57], wrist 0.75 cm [0.29, 1.49]; median [95% confidence interval]. Similar values were found for the other subjects (not reported).

Figure 1A is representative of the arm posture before movement. Specifically, the joint center positions for subject 1 during the 200 ms before movement onset across all task conditions were as follows: jugular notch (11.34 cm, −31.62 cm, −7.3 cm), right shoulder (32.72 cm, −33.23 cm, −6.86 cm), right elbow (27.37 cm, −13.13 cm, −20.27 cm), and right wrist (4.78 cm, −0.11 cm, −10.01 cm). The origin was the handle’s lock position, the +x-direction pointed to the subject’s right, +y-direction pointed ahead of the subject, and +z-direction pointed upward. Similar values were found for the other subjects (not reported).

### EMG patterns co-varied with force threshold and with target zone

We recorded bipolar surface EMG from 8 muscle groups in the arm and shoulder (Figure 6) and found that muscle activity gradually ramped up before movement onset at time 0 (solid vertical line in Figure 6), particularly for task conditions with the highest force threshold. Additionally, EMG activity of most muscle groups was elevated before and after movement onset, suggesting a level of co-activation of antagonist muscle groups.

**Figure 6.**
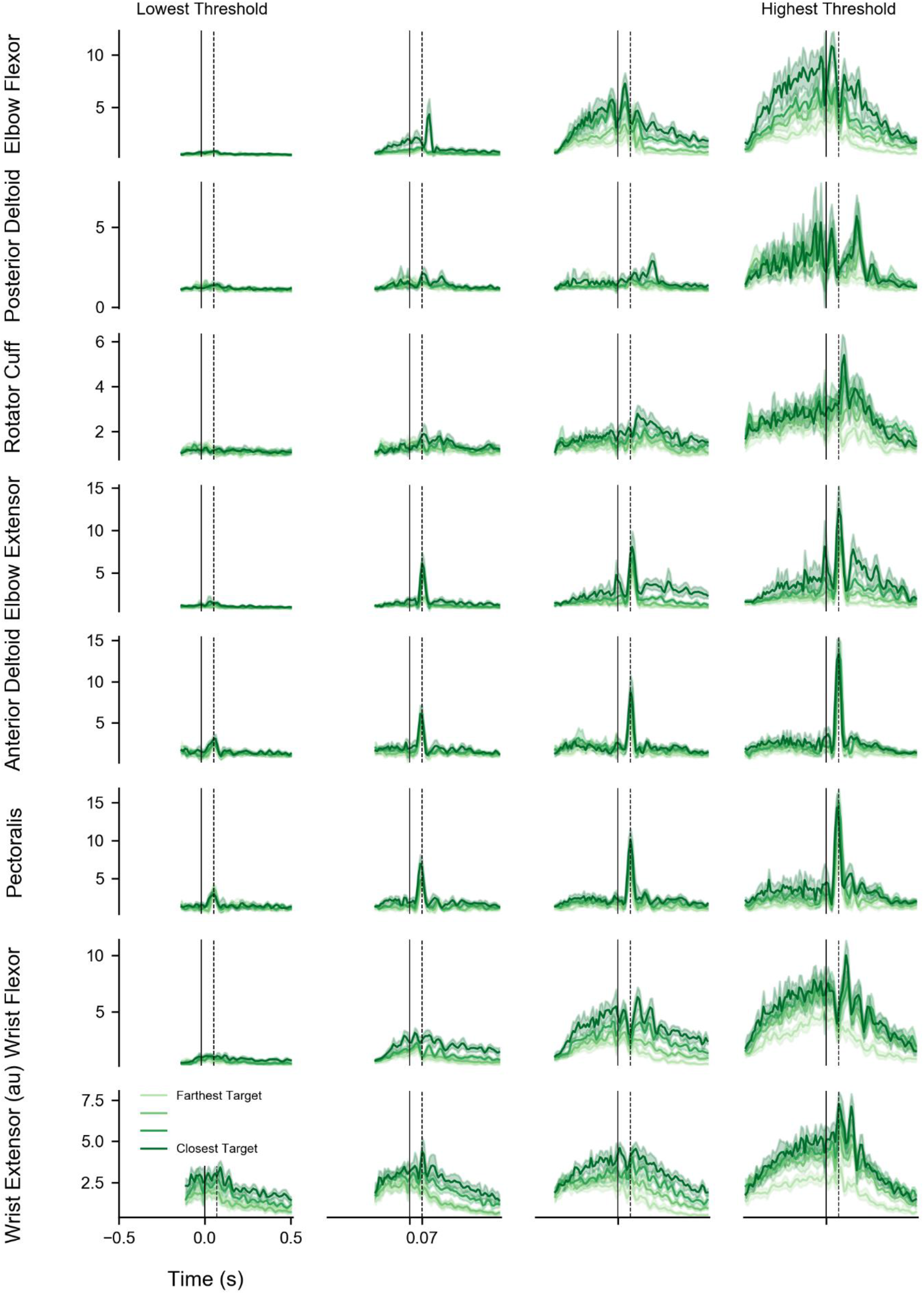
Time-varying EMG across task conditions. The data are from subject 1. Each plot depicts four target zones with the same force threshold. Movement onset was at time 0 (solid vertical line). Some muscles exhibited a strong burst about 70 ms after movement onset (dashed vertical line). The wrist and elbow muscle groups appeared to co-activate, particularly for task conditions that included the highest force threshold. Shading represents the median and 95% confidence interval.

For each muscle and task condition, we found the median and 95% confidence interval of the trial-averaged data. We used a bootstrap to re-calculate the trial average 1000 times. Each calculation used a random sample of the trials, with replacement.

The elbow flexor, posterior deltoid, and rotator cuff were particularly active before movement onset, i.e., while the subject was increasing the force they exerted on the handle. The activity of these muscle groups, and their biomechanical actions, suggest that they might be acting to exert force to cross the force threshold and accelerate the handle along the track.

The elbow extensor, anterior deltoid, and pectoralis activity showed a distinct burst about 70 ms after the handle started to move (dashed vertical line in Figure 6), consistent with a stretch reflex and the functional use of these muscles to decelerate the handle after it started to move.

The wrist flexor and extensor were most consistently modulated across task conditions. One possibility is that the co-activation of these antagonist muscle groups would stiffen the wrist joint and improve the force transfer from the arm to the handle. Another possibility is that this activity may reflect how tightly the subject squeezed the handle, and this may also stiffen the linkage. With our current data, we cannot distinguish between these possibilities.

There are oscillations in the EMG before and after the handle was released. These tend to increase with larger force thresholds (e.g., posterior deltoid). These oscillations and those in the force and acceleration traces are evident in Figures 2A and D and appear to be phase-locked to movement onset. The acceleration pulse occurring at movement onset is in phase with the preceding oscillations, suggesting that they may play a role in overcoming the force threshold.

Comparing a muscle group’s EMG across the force thresholds shows that muscle activity increases with force. In addition, there is a tendency for greater muscle activity for near targets (especially evident at high force thresholds). For a given threshold, the exerted force is similar for different target zones (Figure 2A), despite different EMG patterns, suggesting that the subject was modulating the coactivation of antagonist muscle groups.

### EMG decomposed into potent and null dimensions

We regressed force on EMG during the force ramp and found that a linear model could explain much of the variance (RMSE = 11.68 N +/− 1.44, mean +/− sem across subjects, Supplementary Figure 5). Because there were more dimensions of EMG (8 muscle groups) than dimensions of force (only one along the movement direction), the mapping from EMG to force was redundant. If the mapping from EMG to force was perfectly linear, there would be one direction (i.e., muscle combination) in the multiple-dimensional EMG space that correlated with changes in force. EMG orthogonal to this direction, i.e. in the remaining dimensions of EMG space, would not correlate with force changes.

Using the regression coefficient matrix (equation 2), we found a rotation in the multi-dimensional EMG space that maximized EMG-force correlation in a single dimension. We called EMG projected onto this dimension “potent EMG.” EMG projected onto the remaining 7 dimensions, i.e., dimensions that theoretically did not correlate with changes in force, were summarized using principal component analysis and called “null EMG.”

We found that potent EMG accounted for 21.28% +/− 4.35 of the total variance (mean +/− sem across subjects). The remaining non-potent variance occupied the dimensions that did not correlate as well with changes in force. Null EMG accounted for 77.93% +/− 2.44 of the non-potent variance and 61.71% +/− 4.41 of the total variance (mean +/− sem across subjects).

We designed the null-space analysis under the assumption that balanced changes in the EMG of antagonist muscles produce changes in EMG without concomitant changes in net force. Increases in this type of balanced muscle activation should correspond to changes in stiffness. Although we found that force is generally related to stiffness (Figure 4), stiffness is likely to be better related to null EMG than to potent EMG. We tested this hypothesis by regressing the time-averaged potent and null EMG on stiffness and comparing the slopes, resampling the data 1000 times, with replacement (Supplementary Figure 6). The median values, displayed in figure 7, show that increases in stiffness correspond to greater increases in null EMG compared to potent EMG for 4 out of the 5 subjects (EMG offsets were matched to highlight the slope). This was further confirmed by a histogram of the difference between the null EMG slope and the potent EMG slope (Figure 8).

**Figure 7.**
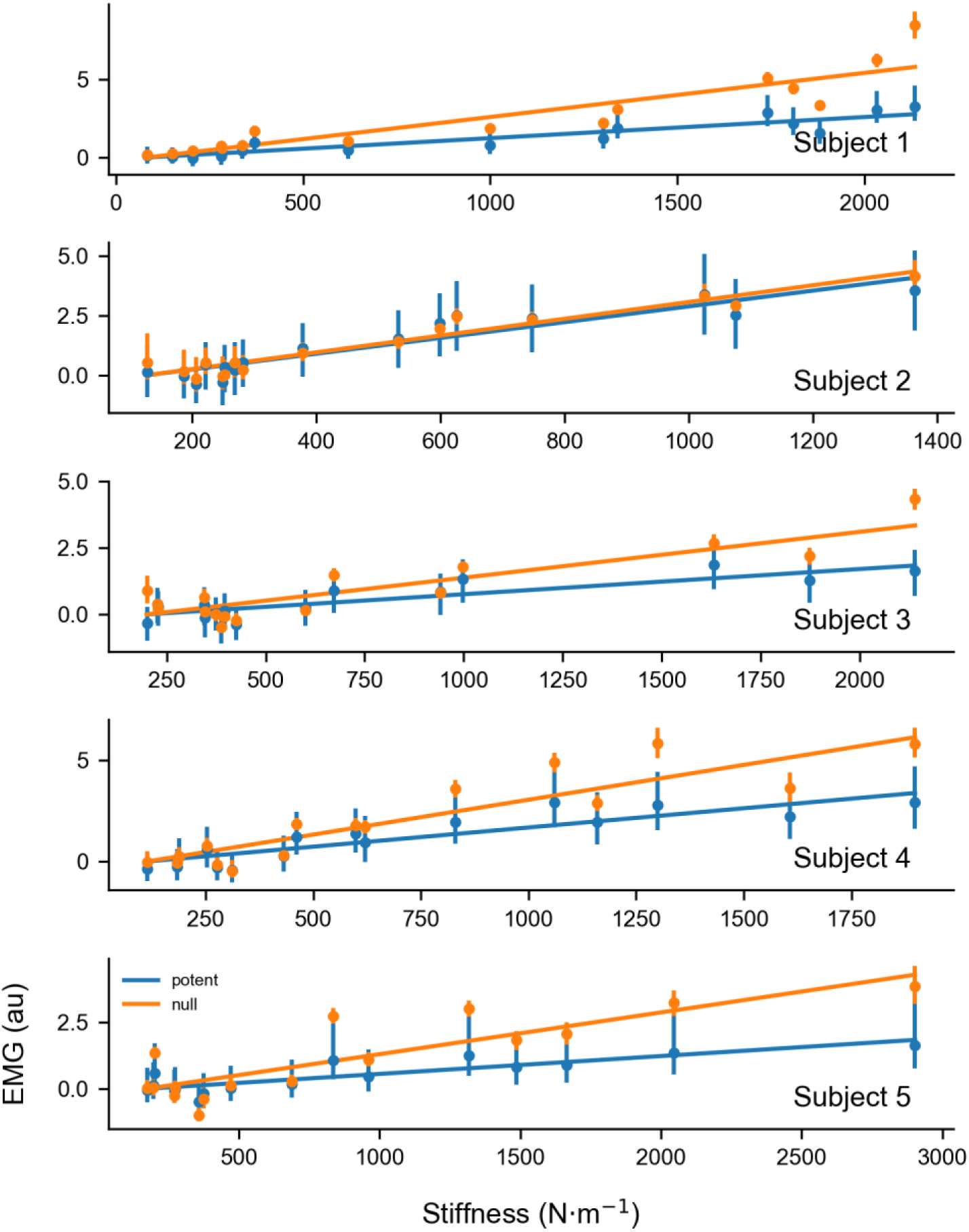
Stiffness has a greater effect on null EMG compared to potent EMG. Each plot depicts the time-averaged EMG for all sixteen conditions. Error bars are the median and 95% confidence interval. EMG offsets were matched to highlight the slope. Subject 1 shows the most distinct difference between null and potent EMG, while subject 2 shows no difference. The potent EMG was the direction in EMG space that was most correlated with force. The null EMG was the orthogonal direction that captured the most variance in the non-potent dimensions.

**Figure 8.**
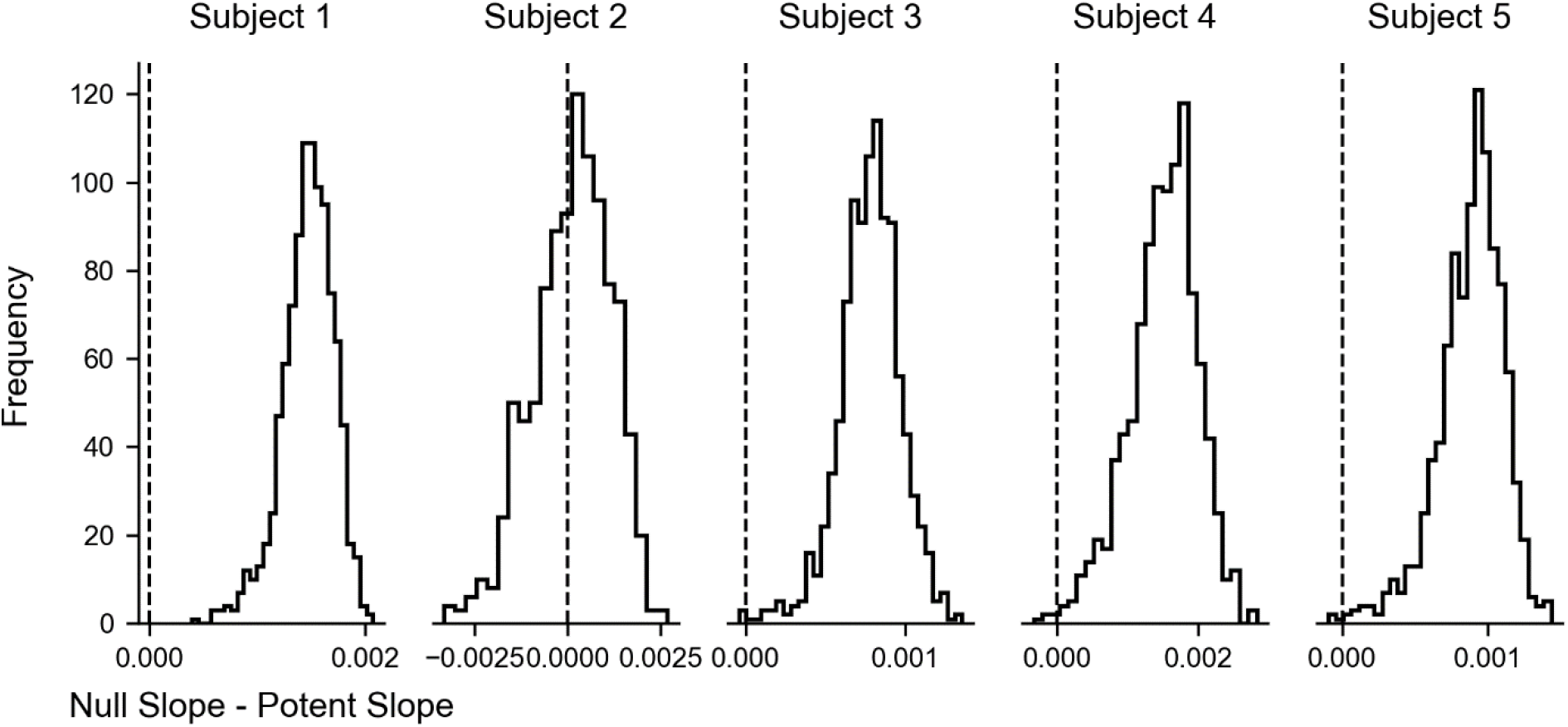
Stiffness has a larger effect on null EMG compared to potent EMG. The slope relating null EMG to stiffness was greater than the slope relating potent EMG to stiffness. The data were resampled 1000 times, with replacement. For each resample, we calculated the difference in slopes relating potent and null EMG to stiffness. Vertical dashed lines represent no difference in the slopes, indicating no difference in the effect of stiffness on potent and null EMG. Positive values indicate that the null EMG slope was greater than the potent EMG slope.

## Discussion

Successfully interacting with an object relies on the ability to coordinate movement of, and the force exerted on, an object. Modulating arm impedance could be an efficient strategy for this type of control. We have designed a ballistic-release paradigm that encourages this strategy while making it possible to dissociate the relations between arm impedance, force, and movement. Of the arm impedance elements studied, stiffness was the only one that varied consistently across task conditions. Other studies have shown that stiffness co-varies with force (McIntyre et al. 1996; Gomi and Osu 1998; Perreault et al. 2002). However, in our task, the linear force-stiffness relation had structure that was target-dependent. Stiffness was related to the target specified before the movement began and was larger for targets that required shorter movements. Our method for relating EMG to stiffness found that separate components of muscle activity varied with force and with stiffness. In designing this study, our aim was to emphasize the use of impedance control as a way of completing a task. These results, from the ballistic-release paradigm, expand upon studies of arm impedance that investigated the isolated effect of force (McIntyre et al. 1996; Gomi and Osu 1998; Perreault et al. 2002) or movement (Gomi and Kawato 1997; Burdet et al. 2001; Darainy et al. 2007; Franklin et al. 2007; Piovesan et al. 2013) and extend the concept that impedance can be specified predictively for object interaction (Lacquaniti et al. 1993; Damm and McIntyre 2008).

### Arm impedance estimation

We found stiffness values that were higher than those reported previously from movements with slower arm speeds and smaller forces (Gomi and Kawato 1997; Darainy et al. 2007). Higher forces and arm speeds are both known to increase stiffness (Latash and Gottlieb 1991; McIntyre et al. 1996). In a study with similar arm speeds, values of stiffness are similar to the values we report (Piovesan et al. 2013). Typically, arm impedance is estimated by perturbing the arm in a way that is external to the task, either by applying a force pulse (Gomi and Kawato 1996, 1997; Piovesan et al. 2013) or a position displacement (Mussa-Ivaldi et al. 1985; Burdet et al. 2001; Darainy et al. 2007; Franklin et al. 2007). In these studies, it is assumed that subjects do not intervene during the perturbation. Arm impedance can be estimated by measuring the force exerted on the object when the arm’s movement is perturbed away from its equilibrium position. However, this method relies on accurately estimating equilibrium positions. In posture maintenance paradigms, position, velocity, and acceleration are zero at this position (Mussa-Ivaldi et al. 1985; Gomi and Osu 1998; Perreault et al. 2002). It is more difficult to estimate impedance during movement (Gomi and Kawato 1997; Darainy et al. 2007). Our task was designed so that the force perturbation, unlocking the handle, was a specific component of the task. Instead of using movement perturbations interspersed throughout the task, we estimated arm impedance using a physical dynamical model (Viviani and Terzuolo 1973; Gomi and Kawato 1997; Burdet et al. 2001; Darainy et al. 2007) based on the initial displacement of the hand as it was released. Impedance, in our model, was calculated with the assumption that the equilibrium position (the target) was constant, and that at this position, velocity and acceleration were both zero (Polit and Bizzi 1979). This assumption was also used in a similar ballistic-release paradigm (Viviani and Terzuolo 1973).

For a given force threshold, the force exerted toward the subject was correlated with stiffness (Supplementary Figure 7). The directional nature of stiffness and force makes it highly unlikely that the off-axis force was causal to the modulation of on-axis stiffness. Instead, a simple correlation can be described (Supplementary Figure 8). Two imaginary muscles could exert force in opposite directions along the track, but in the same direction toward the subject. If the projection of the two muscle forces along the track canceled out, then the on-axis force would be zero and the force exerted toward the subject would be non-zero. Proportional modulation of the muscle forces would increase both the stiffness along the track and the force exerted toward the subject. For this reason, the definition of potent and null EMG also considered only force along the track. Considering force in all three dimensions would have prevented certain behavioral strategies and was not part of the instructions to the subjects.

### Muscle-related stiffness during the ballistic movement

Reaching toward a target is composed of an initial rapid stereotypic movement followed by a homing phase composed of multiple small submovements (Woodworth 1899; Meyer et al. 1988). The initial phase is considered too rapid for concurrent corrections (Elliott et al. 2010) and the arm effectively behaves as a spring-mass-damper system (Viviani and Terzuolo 1973; Hogan 1985b). Upon release, our subjects moved in a way that was similar to this initial reaching phase in that the task constraints encouraged a behavior that was ballistic. This is consistent with a control strategy characterized by preset stiffness and damping.

The first 200 ms following movement onset took place in the absence of corrective interventions to the movement. Since those movements depended on threshold and target, it is likely that the combination of muscle activations were pre-adjusted to reflect the behavioral conditions of the task, as muscle activation contributes to both the force exerted on the handle and to arm impedance (Hogan 1985b).

During this time window, it is likely that feedback-related changes in EMG take place, as evidenced by the EMG response we saw at 70 ms (Figure 6). These responses may be comprised of spinal and cortical reflexes and could contribute to both stiffness and damping. These responses are known to change with task requirements (Kurtzer et al. 2009; Dimitriou et al. 2013; Pruszynski et al. 2014) and could be preset by the nervous system to control the arm during the movement. Our analysis shows that, by modeling stiffness in a window that incorporates these responses, we could obtain a match between the equilibrium point and arrest position, suggesting that the impedance-control framework may incorporate these responses.

### Impedance control

Results from a similar paradigm suggest that the equilibrium position is set before the force begins to increase (Elliott et al. 1999). In that study, the handle was unlocked in random catch trials before or after the force threshold was crossed. Since force increased continuously before the handle was released, a pure force-dependent strategy would result in shorter displacements for earlier releases. However, the handle’s displacement was not related to the time of release in the catch trials, showing that a simple predictive force strategy was probably not used in the task. Furthermore, force at high magnitudes is susceptible to signal-dependent noise (Harris and Wolpert 1998; Franklin et al. 2004), making it difficult, as in our task, to predict when force would cross the threshold and unlock the handle, again arguing against the idea of a scheme relying only on predictive force control.

Although our results suggest that arm impedance is set before the movement takes place, in theory, subjects could perform the ballistic-release paradigm without changing arm impedance. If a subject could predict when the handle would be unlocked, it would not be necessary to change arm impedance for different targets. Instead, a set of muscles could be activated to generate the force needed to unlock the handle, followed by the activation of a different set that would generate the precise force needed to decelerate the handle to stop in the target. In contrast to co-activation, this type of control would be energetically efficient (Franklin et al. 2004) and could be implemented using implicit and/or explicit knowledge of the physical plant’s mechanics (i.e., mapping activation to force) along with the object’s properties (i.e. the force needed to unlock the handle). The subject’s internal model would encompass this knowledge and could be used to precisely control the transition from isometric force to movement control. Although this would be energetically efficient, precise timing would be required. The additional details needed for this scheme would increase information loading in the system and could result in slower movement.

Our results suggest that subjects chose to control both force and movement together via impedance control, an idea consistent with other studies which found impedance control to be preferred when the relation between force and movement is uncertain (Thoroughman and Shadmehr 1999; Takahashi et al. 2001; Franklin et al. 2003). These concepts also fit under the umbrella of the equilibrium point hypothesis (Feldman 1966, 1986) and its extensions (Bizzi et al. 1984; Hogan 1985b; Flash 1987). In this framework, the force exerted on the handle in our task is controlled only indirectly. It depends on the desired movement, the actual movement perceived from sensory feedback, and the preset arm impedance.

## Conclusion

This ballistic-release paradigm used in this task encouraged subjects to adopt a strategy in which they simultaneously activated their muscles, creating a virtual spring that arrested a fast arm movement in a specified target. Subjects adjusted their arm impedance to achieve the desired displacement for each combination of force threshold and target position. By modeling the arm’s impedance in the short interval following release, we found that arm stiffness had changed in a way that anticipated the displacement needed to reach the target. We assessed the relation between arm impedance and muscle activation, finding EMG patterns that were less correlated with changes in force. This “null” component was, instead, highly correlated with stiffness, suggesting that subjects used their muscles to modulate arm impedance without changing force. The ability to separate changes of force, position and stiffness and how these are associated with different components of muscle activation suggests that anticipatory changes in impedance may be a cardinal feature of manipulative control. Because this paradigm demonstrates this aspect of control, it will be useful for studying the relation between cortical neural activity and arm impedance (Humphrey and Reed 1983). Future enhancement of this paradigm to include multiple directions (Darainy et al. 2007) and time-varying estimates of arm impedance (Lacquaniti et al. 1993; Piovesan et al. 2013) will make it possible to generalize the control of object interaction to a wider range of behaviors.

## Supplementary Material

**Supplementary Figure 1.**
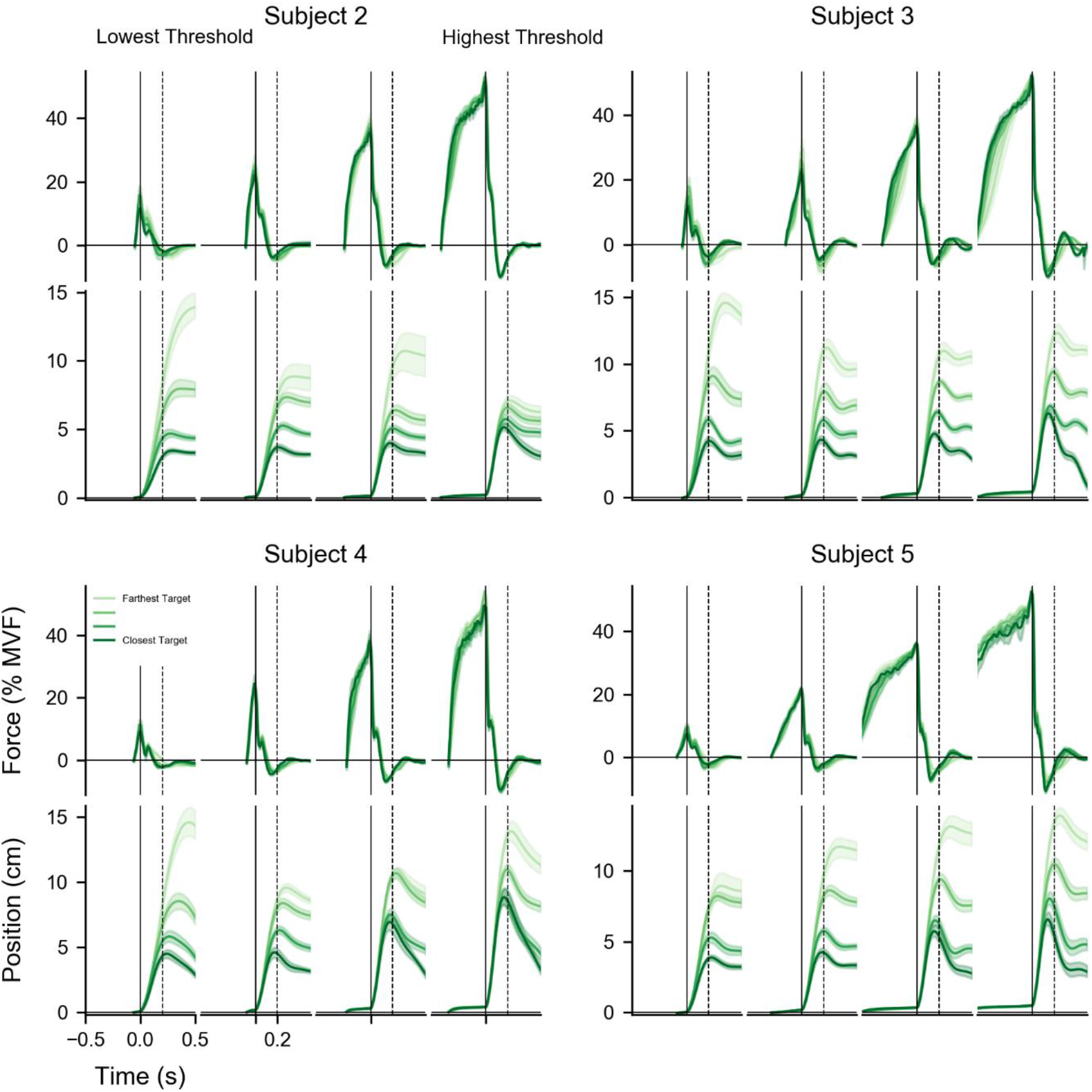
Behavioral data for all subjects. Movement varied considerably across subjects. However, across-trial variability was much less. For some conditions, the subjects returned the handle to the start position without holding the handle in place (see Subject 4, highest threshold). Additionally, Subject 2 adopted the strategy for the highest threshold of arresting the handle in the same position, regardless of the target. The data are trial-averaged and shading represents the median and 95% confidence interval. Each plot depicts four target zones with the same force threshold. Movement onset began at time 0 (solid vertical line). The maximum voluntary force (MVF) for Subject 2-5 was 100, 160, 200, and 200 N.

**Supplementary Figure 2.**
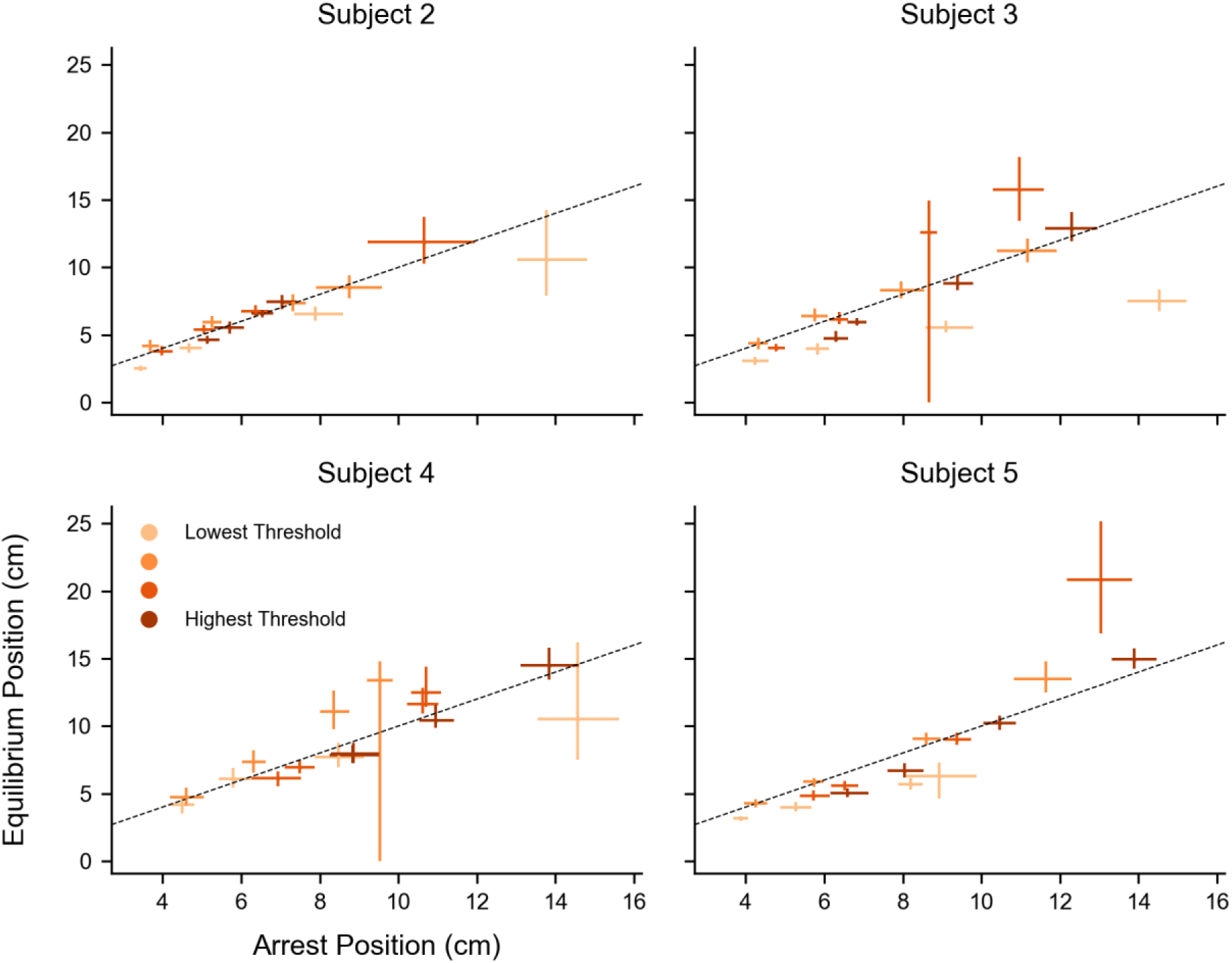
Equilibrium position for all subjects. The distance to the equilibrium position (EP) was similar to the distance to the arrest position, suggesting that the physical dynamical model could describe important aspects of the behavior. The distance to the EP was generally a little less than the distance to the arrest position, especially for Subject 5, which could be explained by an underdamped physical system and result in the position overshoot observed in Supplementary Figure 1. Markers are the median and 95% confidence interval.

**Supplementary Figure 3.**
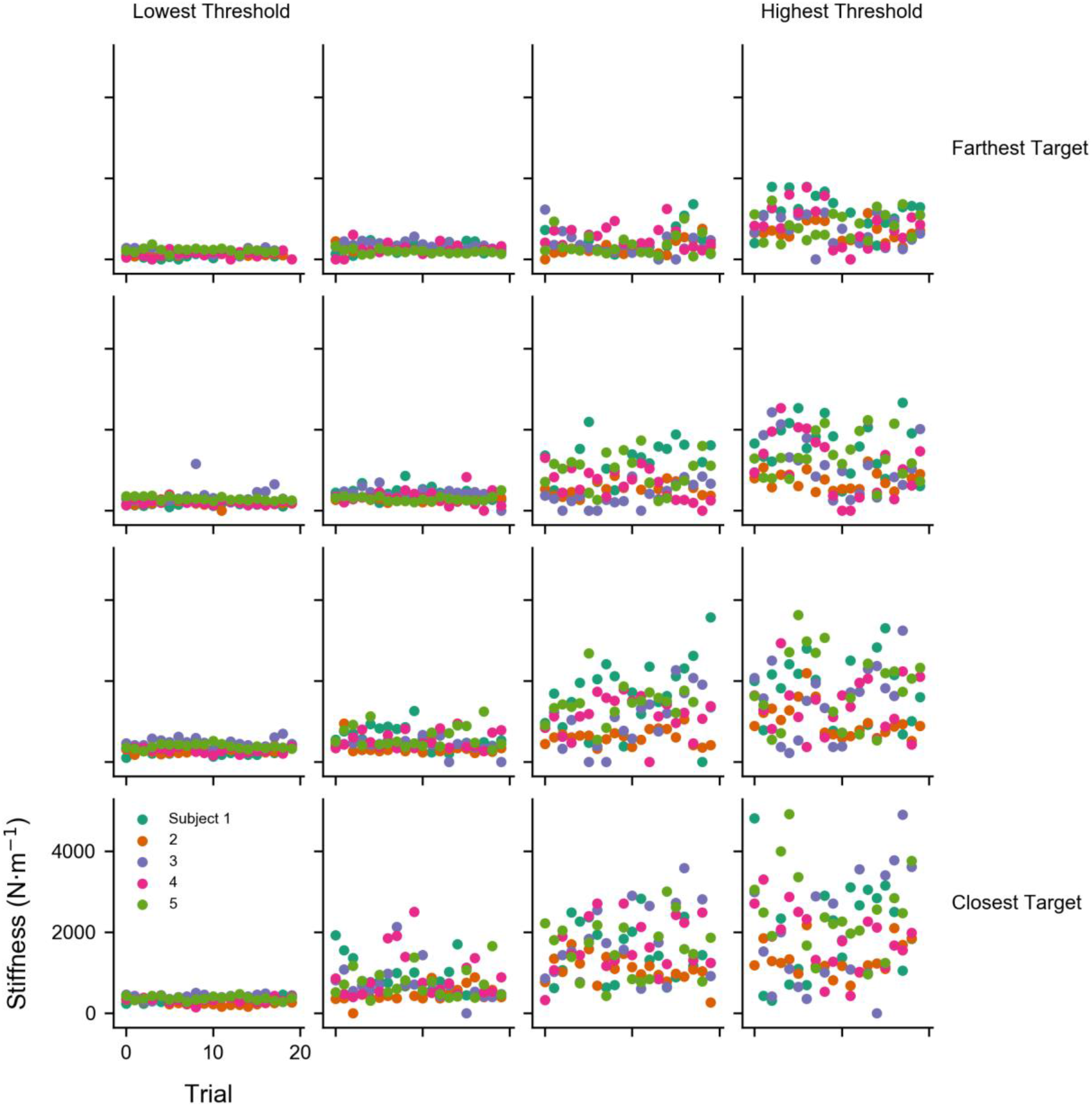
Stiffness across trials for all subjects and task conditions. Subjects did not appear to modulate their stiffness consistently across trials for a given condition. However, variability within a given condition did increase with stiffness, suggesting signal-dependent noise that could be linked to similar signal-dependent noise in muscle force.

**Supplementary Figure 4.**
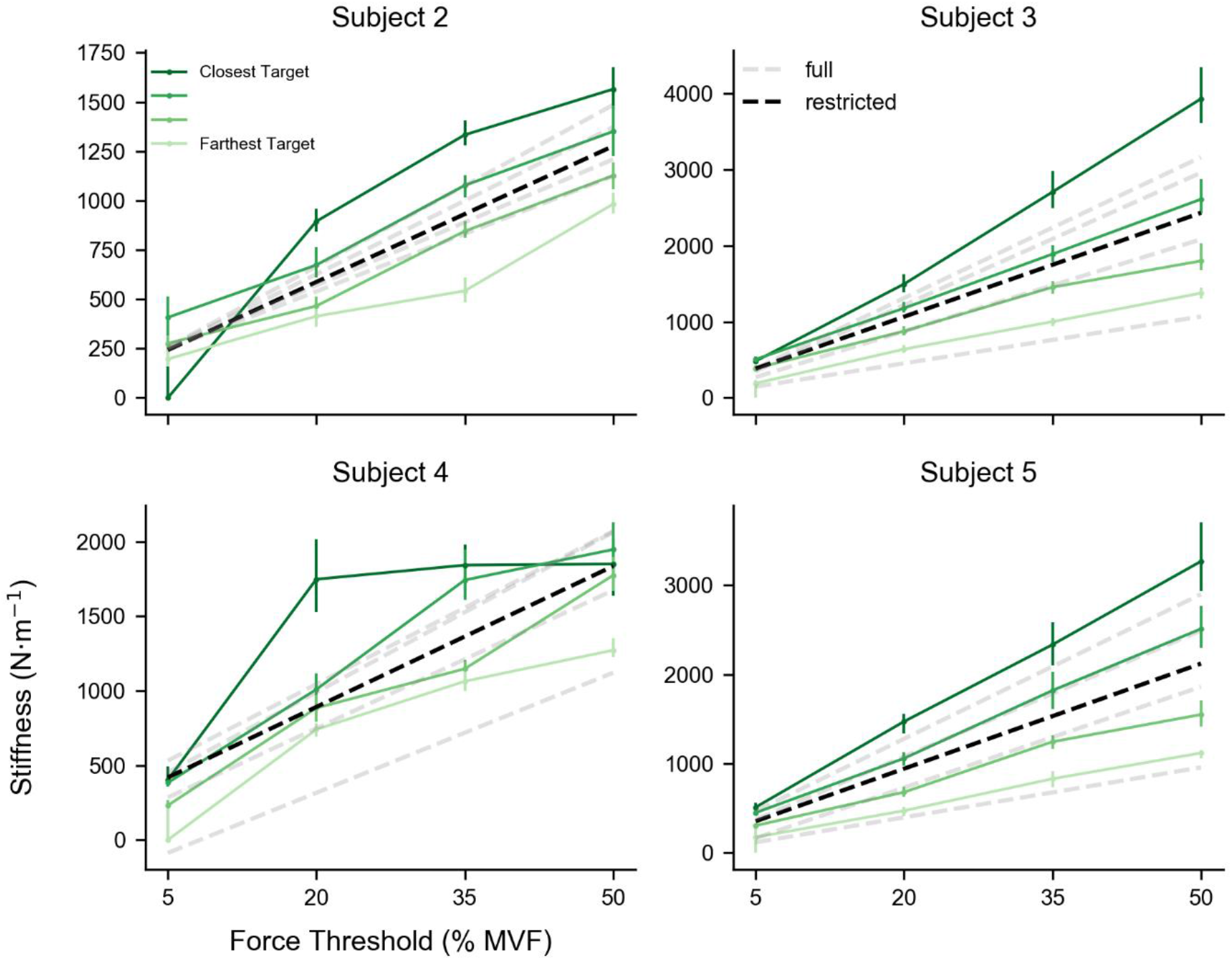
Stiffness depended on both force and movement. Stiffness exhibited a strong linear relation with force threshold, particularly for Subjects 3 and 5. However, the slope and offset of the linear relation depended on the target. Error bars indicate median and 95% confidence interval. The dashed grey lines represent the linear fit of the full regression of stiffness on force threshold and arrest position. The dashed black line represents the linear fit of the restricted regression of stiffness on force threshold alone. The physical dynamical model was fit using 200 ms of data beginning at movement onset.

**Supplementary Figure 5.**
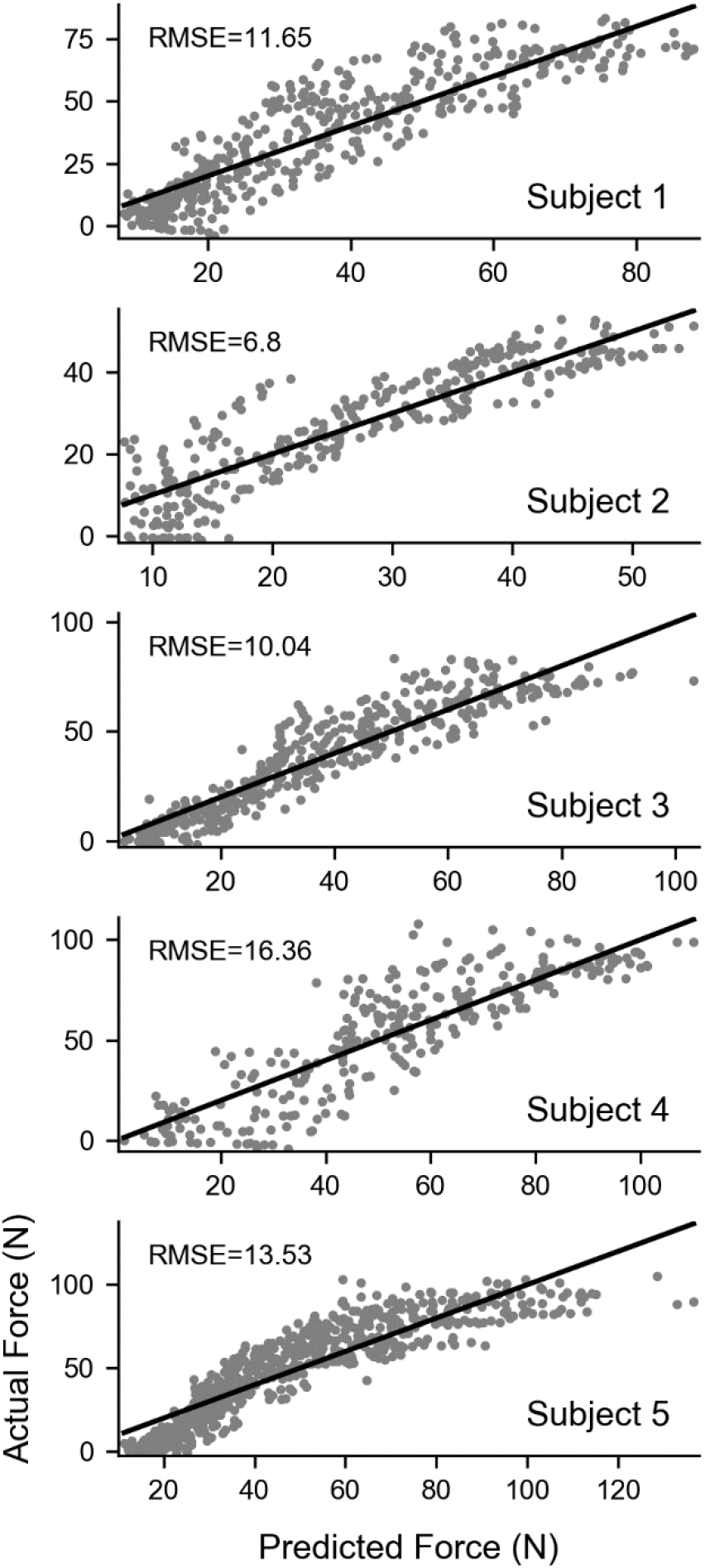
EMG approximates force. Time-varying force during the force ramp was regressed on the EMG from all 8 muscles. The linear model was able to consistently capture variability across all subjects. The data displayed are the median predicted and actual values from the bootstrap. The dark black line is the unity line representing a perfect fit. Although the small deviations from the unity line cold be due to the non-linear relation between EMG and force, a linear model is a good approximation.

**Supplementary Figure 6.**
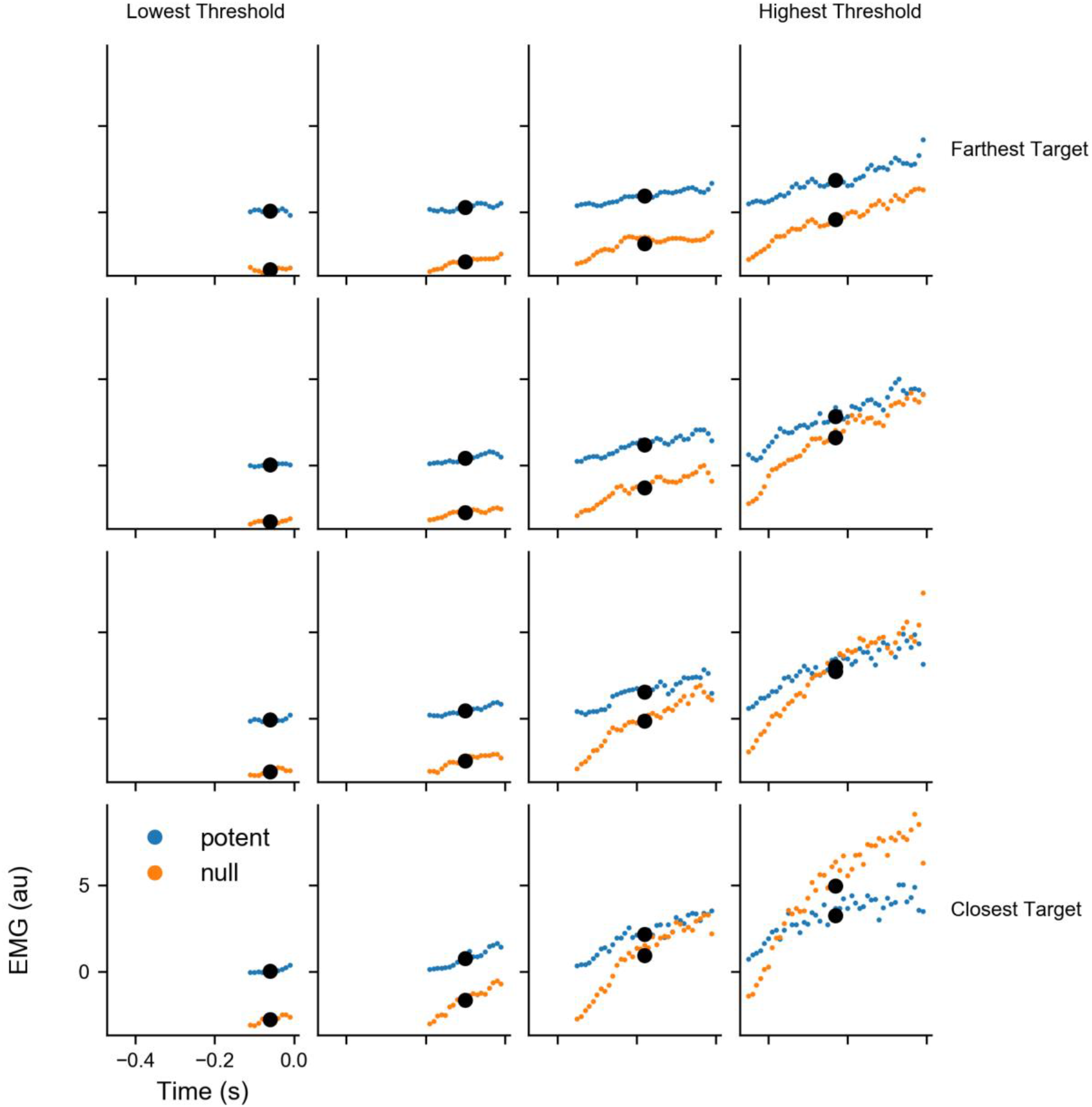
Example of time-averaged EMG for subject 1. There was a single value of stiffness for each task condition. To determine the effect of stiffness on potent and null EMG, we averaged EMG across time before movement onset. The result was a single potent and null EMG value for each task condition that was first averaged across trials and then averaged across time. The confidence intervals were calculated by resampling 1000 times from the 20 trials per condition. The colored data are the time series EMG for a single sample and the large black dots are the averages.

**Supplementary Figure 7.**
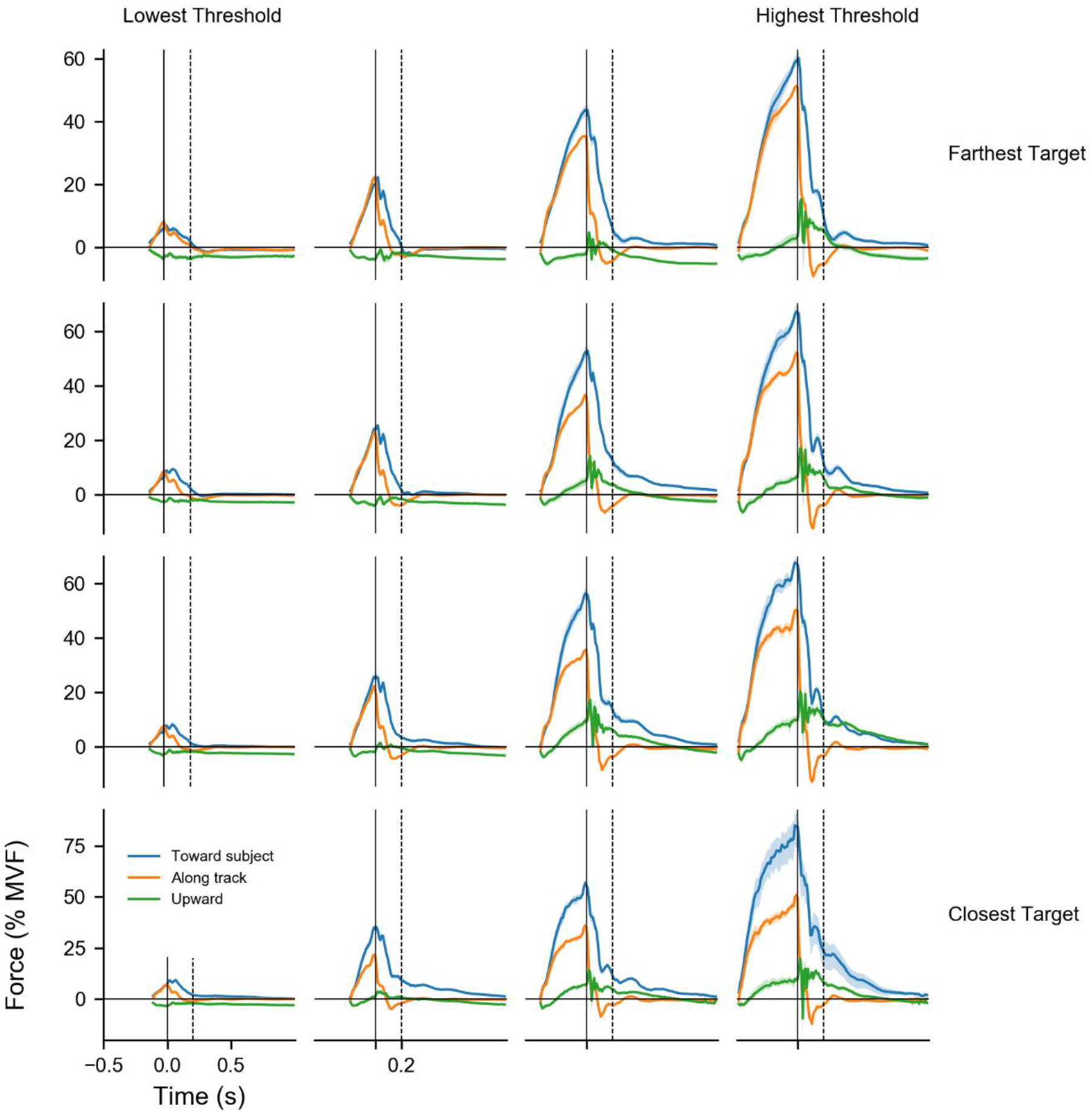
Force in all 3 dimensions. Although force exerted along the track did not vary across conditions with the same force threshold, force exerted toward the subject increased as the target moved closer to the start position. The correlation between the force exerted toward the subject and stiffness was likely due to the biomechanics of the arm, where modulation of off-axis muscle force enabled modulation of on-axis co-activation.

**Supplementary Figure 8.**
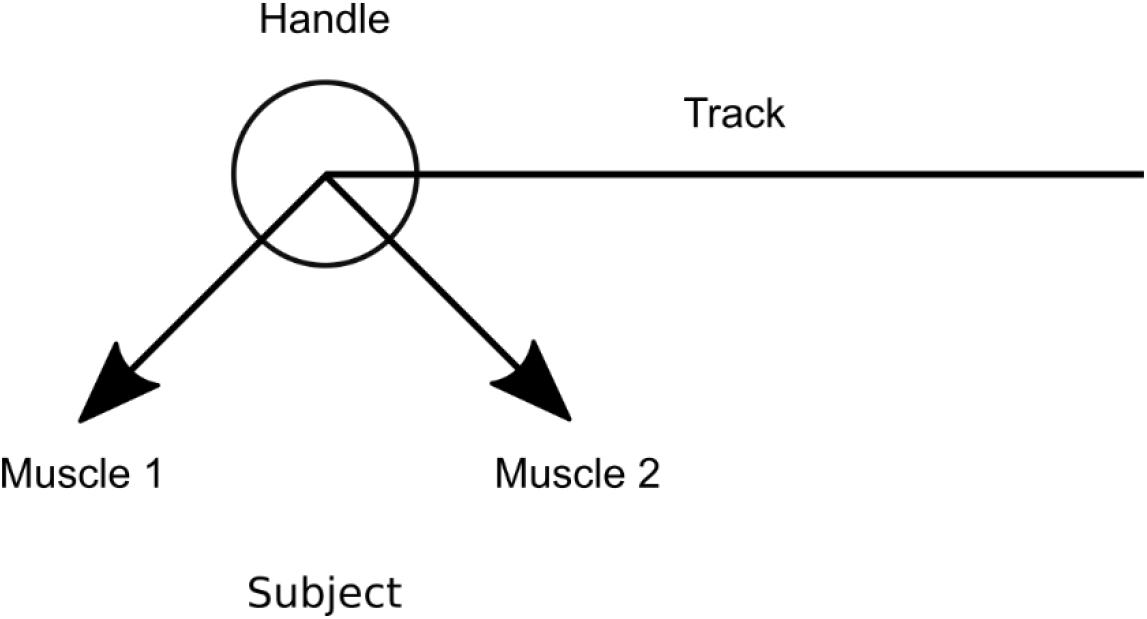
Off-axis forces could correlate with on-axis stiffness. Two imaginary muscles exert force on the handle according to the labeled vectors. If the magnitude of the muscle forces are equal, then the sum of the two muscle forces would result in zero force along the track and a non-zero force toward the subject. In addition, increasing the muscle force would increase both the force toward the subject and the stiffness along the track, without changing the force along the track.

## Acknowledgements

Supported by a Google Faculty Award to A.B.S. The authors would like to thank Steve Chase, Josue Orellana, Neville Hogan, Jordan Williams, and Rex Tien for helpful comments.

## References

Bizzi E, Accornero N, Chapple W, Hogan N. Arm trajectory formation in monkeys. Exp Brain Res 46: 139–143, 1982.

Bizzi E, Accornero N, Chapple W, Hogan N. Posture control and trajectory formation during arm movement [Online]. J Neurosci 4: 2738–44, 1984. http://www.ncbi.nlm.nih.gov/pubmed/6502202.

Burdet E, Osu R, Franklin DW, Milner TE, Kawato M. The central nervous system stabilizes unstable dynamics by learning optimal impedance. Nature 414: 446–449, 2001.

Crevecoeur F, Scott SH. Beyond Muscles Stiffness: Importance of State-Estimation to Account for Very Fast Motor Corrections. PLoS Comput Biol 10, 2014.

Damm L, McIntyre J. Physiological basis of limb-impedance modulation during free and constrained movements. J Neurophysiol 100: 2577–2588, 2008.

Darainy M, Towhidkhah F, Ostry DJ. Control of hand impedance under static conditions and during reaching movement. J Neurophysiol 97: 2676–85, 2007.

Dimitriou M, Wolpert DM, Franklin DW. The Temporal Evolution of Feedback Gains Rapidly Update to Task Demands. J Neurosci 33: 10898–10909, 2013.

Elliott D, Hansen S, Grierson LEM, Lyons J, Bennett SJ, Hayes SJ. Goal-Directed Aiming: Two Components but Multiple Processes. Psychol Bull 136: 1023–1044, 2010.

Elliott D, Heath M, Binsted G, Ricker KL, Roy EA, Chua R. Goal-Directed Aiming: Correcting a Force-Specification Error With the Right and Left Hands. J Mot Behav 31: 309–324, 1999.

Feldman AG. On the functional tuning of the nervous system in movement control or preservation of stationary pose. II. Adjustable parameters in muscles [Online]. Biofizika 11: 498–508, 1966. http://www.ncbi.nlm.nih.gov/pubmed/5999810.

Feldman AG. Once more on the equilibrium-point hypothesis (lambda model) for motor control [Online]. J Mot Behav 18: 17–54, 1986. http://www.ncbi.nlm.nih.gov/pubmed/15136283.

Flanagan JR, Bowman MC, Johansson RS. Control strategies in object manipulation tasks. Curr Opin Neurobiol 16: 650–659, 2006.

Flash T. The control of hand equilibrium trajectories in multi-joint arm movements. Biol Cybern 57: 257–274, 1987.

Flash T, Hogan N. The coordination of arm movements: an experimentally confirmed mathematical model [Online]. J Neurosci 5: 1688–1703, 1985. http://www.ncbi.nlm.nih.gov/pubmed/4020415.

Franklin DW, Liaw G, Milner TE, Osu R, Burdet E, Kawato M. Endpoint Stiffness of the Arm Is Directionally Tuned to Instability in the Environment. J Neurosci 27: 7705–7716, 2007.

Franklin DW, Osu R, Burdet E, Kawato M, Milner TE. Adaptation to Stable and Unstable Dynamics Achieved By Combined Impedance Control and Inverse Dynamics Model. J Neurophysiol 90: 3270–3282, 2003.

Franklin DW, So U, Kawato M, Milner TE. Impedance control balances stability with metabolically costly muscle activation. J Neurophysiol 92: 3097–105, 2004.

Franklin DW, Wolpert DM. Computational Mechanisms of Sensorimotor Control. Neuron 72: 425–442, 2011.

Gomi H, Kawato. Equilibrium-point control hypothesis examined by measured arm stiffness during multijoint movement. Science (80-) 272: 117–20, 1996.

Gomi H, Kawato M. Human arm stiffness and equilibrium-point trajectory during multi-joint movement. Biol Cybern 76: 163–171, 1997.

Gomi H, Osu R. Task-dependent viscoelasticity of human multijoint arm and its spatial characteristics for interaction with environments [Online]. J Neurosci 18: 8965–8978, 1998. http://www.ncbi.nlm.nih.gov/pubmed/9787002.

Gribble PL, Mullin LI, Cothros N, Mattar A. Role of Cocontraction in Arm Movement Accuracy. J Neurophysiol 89: 2396–2405, 2003.

Harris CM, Wolpert DM. Signal-dependent noise determines motor planning. Nature 394: 780–4, 1998.

Hill A V. The series elastic component of muscle [Online]. Proc R Soc London Ser B, Biol Sci 137: 273–280, 1950. http://www.ncbi.nlm.nih.gov/pubmed/15430325.

Hogan N. An organizing principle for a class of voluntary movements [Online]. J Neurosci 4: 2745–2754, 1984a. http://www.ncbi.nlm.nih.gov/pubmed/6502203.

Hogan N. Impedance Control: An Approach to Manipulation. In: IEEE American Control Conference. 1984b, p. 304–313.

Hogan N. Impedance Control: An Approach to Manipulation: Part II - Implementation. J Dyn Syst Meas Control 107: 1–7, 1985a.

Hogan N. The Mechanics of Multi-Joint Posture and Movement Control. Biol Cybern 331: 315–331, 1985b.

Humphrey DR, Reed DJ. Separate cortical systems for control of joint movement and joint stiffness: reciprocal activation and coactivation of antagonist muscles [Online]. Adv Neurol 39: 347–72, 1983. http://www.ncbi.nlm.nih.gov/pubmed/6419553.

Joyce GC, Rack PM. Isotonic lengthening and shortening movements of cat soleus muscle [Online]. J Physiol 204: 475–491, 1969. http://jp.physoc.org/content/204/2/475.abstract.

Kadiallah A, Liaw G, Kawato M, Franklin DW, Burdet E. Impedance control is selectively tuned to multiple directions of movement. J Neurophysiol 106: 2737–2748, 2011.

Kalaska JF, Crammond DJ. Cerebral cortical mechanisms of reaching movements. Science (80-) 255: 1517–1523, 1992.

Kaufman MT, Churchland MM, Ryu SI, Shenoy KV. Cortical activity in the null space: permitting preparation without movement. Nat Neurosci 17: 440–8, 2014.

Kawato M. Internal models for motor control and trajectory planning. Curr Opin Neurobiol 9: 718–727, 1999.

Kurtzer I, Pruszynski JA, Scott SH. Long-Latency Responses During Reaching Account for the Mechanical Interaction Between the Shoulder and Elbow Joints. J Neurophysiol 102: 3004–3015, 2009.

Lacquaniti F, Carrozzo M, Borghese NA. Time-varying mechanical behavior of multijointed arm in man. J Neurophysiol 69: 1443–1464, 1993.

Lacquaniti F, Licata F, Soechting J. The Mechanical Behavior of the Human Forearm in Response to Transient Perturbations. Biol. Cybern.

Latash ML, Gottlieb GL. Reconstruction of shifting elbow joint compliant characteristics during fast and slow movements. Neuroscience 43: 697–712, 1991.

McIntyre J, Mussa-Ivaldi F, Bizzi E. The control of stable postures in the multijoint arm. Exp Brain Res 110: 248–264, 1996.

Meyer DE, Abrams RA, Kornblum S, Wright CE, Smith JE. Optimality in human motor performance: ideal control of rapid aimed movements [Online]. Psychol Rev 95: 340–370, 1988. http://www.ncbi.nlm.nih.gov/pubmed/3406245.

Milner TE, Franklin DW. Impedance control and internal model use during the initial stage of adaptation to novel dynamics in humans. J Physiol 567: 651–664, 2005a.

Milner TE, Franklin DW. Impedance control and internal model use during the initial stage of adaptation to novel dynamics in humans. J Physiol 567: 651–664, 2005b.

Mussa-Ivaldi FA, Hogan N, Bizzi E. Neural, Mechanical, and Geometric Factors Subserving Arm Posture in Humans. J Neurosci 5: 2732–2743, 1985.

Nichols TR, Houk JC. Improvement in linearity and regulation of stiffness that results from actions of stretch reflex. J Neurophysiol 39: 119–142, 1976.

Osu R, Burdet E, Franklin DW, Milner TE, Kawato M. Different Mechanisms Involved in Adaptation to Stable and Unstable Dynamics. J Neurophysiol 90: 3255–3269, 2003.

Osu R, Gomi H. Multijoint Muscle Regulation Mechanisms Examined by Measured Human Arm Stiffness and EMG Signals. J Neurophysiol 81: 1458–1468, 1999.

Perreault EJ, Kirsch RF, Crago PE. Voluntary Control of Static Endpoint Stiffness During Force Regulation Tasks. J Neurophysiol 87: 2808–2816, 2002.

Piovesan D, Pierobon A, DiZio P, Lackner JR. Experimental measure of arm stiffness during single reaching movements with a time-frequency analysis. J Neurophysiol 110: 2484–2496, 2013.

Polit A, Bizzi E. Characteristics of motor programs underlying arm movements in monkeys [Online]. J Neurophysiol 42: 183–94, 1979. http://www.ncbi.nlm.nih.gov/pubmed/107279.

Pruszynski JA, Kurtzer I, Lillicrap TP, Scott SH. Temporal Evolution of Automatic Gain-Scaling. J Neurophysiol 102: 992–1003, 2009.

Pruszynski JA, Omrani M, Scott SH. Goal-Dependent Modulation of Fast Feedback Responses in Primary Motor Cortex. J Neurosci 34: 4608–4617, 2014.

Rack PMH, Westbury DR. The short range stiffness of active mammalian muscle and its effect on mechanical properties. J Physiol 240: 331–350, 1974.

Rancourt D, Hogan N. Stability in Force-Production Tasks. J Mot Behav 33: 193–204, 2001.

Scott SH. Optimal feedback control and the neural basis of volitional motor control. Nat Rev Neurosci 5: 532–545, 2004.

Scott SH, Cluff T, Lowrey CR, Takei T. Feedback control during voluntary motor actions. Curr Opin Neurobiol 33: 85–94, 2015.

Selen LPJ, Beek PJ, van Dieën JH. Impedance is modulated to meet accuracy demands during goal-directed arm movements. Exp Brain Res 172: 129–138, 2006.

Shadmehr R, Mussa-Ivaldi FA. Adaptive representation of dynamics during learning of a motor task. J Neurosci 14: 3208–24, 1994.

Takahashi CD, Scheidt RA, Reinkensmeyer DJ. Impedance Control and Internal Model Formation When Reaching in a Randomly Varying Dynamical Environment. J Neurophysiol 86: 1047–1051, 2001.

Thoroughman KA, Shadmehr R. Electromyographic correlates of learning an internal model of reaching movements [Online]. J Neurosci 19: 8573–8588, 1999. http://www.ncbi.nlm.nih.gov/pubmed/10493757.

Trumbower RD, Krutky MA, Yang B-S, Perreault EJ. Use of Self-Selected Postures to Regulate Multi-Joint Stiffness During Unconstrained Tasks. PLoS One 4: e5411, 2009.

Viviani P, Terzuolo CA. Modeling of a simple motor task in man: intentional arrest of an ongoing movement [Online]. Kybernetik 14: 35–62, 1973. http://www.ncbi.nlm.nih.gov/pubmed/4777328.

Wolpert DM, Ghahramani Z. Computational principles of movement neuroscience. Nat Neurosci 3: 1212–1217, 2000.

Woodworth RS. Accuracy of voluntary movement. Psychol Rev Monogr Suppl 3: i–114, 1899.

